# Comparative transcriptomic analysis of *Zymoseptoria tritici* reveals interaction-specific gene expression patterns during susceptible, resistant, and non-host interactions

**DOI:** 10.1101/2023.11.20.567875

**Authors:** Sandra V. Gomez-Gutierrez, Cassidy R. Million, Namrata Jaiswal, Michael Gribskov, Matthew Helm, Stephen B. Goodwin

## Abstract

*Zymoseptoria tritici* is responsible for Septoria tritici blotch, a disease causing significant annual yield losses in wheat. To investigate infection phase-specific gene expression in the pathogen, we analyzed gene expression during infection of susceptible (Taichung 29) and resistant (Veranopolis and Israel 493) wheat cultivars, plus the non-host species barley at 1, 3, 6, 10, 17 and 23 days post inoculation (DPI). There were dramatic differences in pathogen gene expression at 10 DPI in the susceptible compared to both resistant interactions. The most pronounced differences in pathogen gene expression were observed at 3 DPI in both the susceptible and resistant host interactions compared to the non-host interaction. Thirty-one putative effectors showed early expression during the susceptible interaction compared to the non-host interaction, and six effectors were selected for subcellular localization studies. Using *Agrobacterium*-mediated transient expression in *Nicotiana benthamiana*, subcellular localization assays revealed that two candidate effectors, Mycgr3109710 and Mycgr394290, localized to putative mobile cytosolic bodies when expressed without their signal peptides, suggesting potential roles in intracellular signaling or host gene regulation. When expressed with their native signal peptides, four candidate effectors localized to the cytosol, while one effector, Mycgr3107904, did not accumulate to detectable levels, as shown by immunoblot analysis, indicating degradation. Comparison of pathogen gene expression in the susceptible host to expression in the resistant hosts, allowed us to identify genes that are expressed during the transition from biotrophic to necrotrophic growth at 10 DPI. Comparison of pathogen gene expression in resistant and susceptible hosts, versus in the non-host barley, allowed us to identify genes involved in initial colonization and host recognition. In addition, our study contributes to understanding candidate effectors that are activated early during infection and may play a role in the initial suppression of plant immunity, making them strong candidates for functional characterization.

---

*Zymoseptoria tritici* is a fungal pathogen in the class Dothideomycetes that causes Septoria tritici blotch (STB) disease of bread (*Triticum aestivum*) and durum (*T. turgidum* subsp. *durum*) wheat (Kettles & Kanyuka, 2016; Palma-Guerrero et al., 2017). STB is one of the most yield-limiting diseases of wheat globally and may result in fifty percent yield loss, especially in regions with high humidity and mild temperatures (Mekonnen et al., 2020; Yang et al., 2022). *Z. tritici* is a hemibiotroph that transitions from biotrophic to necrotrophic growth after a prolonged 7- to 10-day latent period (Ponomarenko et al., 2011; Rudd et al., 2015). During the biotrophic growth phase, the pathogen circumvents recognition through suppression of host immune responses, likely by translocating virulence (effector) proteins. During the necrotrophic phase, the fungus degrades host tissue and presumably extracts nutrients from host plant cells (Marshall et al., 2011; Steinberg, 2015). The hemibiotrophic lifestyle of *Z. tritici* coupled with the availability of a complete, chromosome-level genome (Goodwin et al., 2011) makes this fungal pathogen an attractive model to investigate infection phase-specific gene expression.

Though *Z. tritici* is a hemibiotroph, our knowledge of its infection strategy and lifestyle transition remains limited. Nevertheless, comprehensive RNA sequencing analyses have provided insights into host and pathogen gene expression during a susceptible interaction (Rudd et al., 2015; Yang et al., 2013), transcriptome signatures associated with blastospore and mycelial growth (Francisco et al., 2019), as well as differences in gene expression among *Z. tritici* strains (Palma-Guerrero et al., 2017). Studies have also assessed plant gene expression in resistant-host (incompatible) and non-host interactions (Orton et al., 2017; Reilly et al., 2021), and one study compared the pathogen gene expression profile during infection in a susceptible host (*T. aestivum* L.) and a non-host (*Brachypodium distachyon*) (Kellner et al., 2014). Kellner et al. (2014) found that many *Z. tritici* genes that are up-regulated in wheat encode proteins with putative secretion signals and therefore may play a functional role in host invasion. The most highly differentially expressed genes (DEGs) in *Z. tritici* in the comparison of the susceptible and the non-host interactions showed signatures of positive selection and can be considered candidates for potential roles in host specialization of this pathogen (Kellner et al., 2014).

Differential responses of pathogens to susceptible, resistant, and non-host species are often determined by the molecular mechanisms underlying the interactions, as well as virulence (effector) proteins (Selin et al., 2016). Pathogens often secrete effectors, in part, to circumvent host innate immune responses (Petit-Houdenot & Fudal, 2017; Selin et al., 2016). Once translocated to the host apoplast or cytosol, effector proteins interfere with cell surface-mediated and intracellular-mediated immune responses, thereby promoting pathogenesis and disease development. In *Z. tritici*, some effector proteins have been functionally characterized (Motteram et al., 2009; Marshall et al., 2011; Mirzadi Gohari et al., 2015; Rudd et al., 2015; Ben M’barek et al., 2015; Karki et al., 2021; Thynne et al., 2024). For example, the *Z. tritici* effector Mg3LysM is expressed during early disease development and suppresses chitin-induced defense responses in wheat (Marshall et al., 2011; Rudd et al., 2015). Furthermore, Karki et al. (2021) recently identified an effector from *Z. tritici*, ZtSSP2, that is expressed at 2 days post inoculation (DPI) and interacts with a wheat E3 ubiquitin ligase (TaE3UBQ). Intriguingly, virus-induced gene silencing of TaE3UBQ resulted in increased *Z. tritici* susceptibility, suggesting its encoded protein may function as a regulator of wheat defense responses, and that ZtSSP2 may suppress TaE3UBQ ligase activity to promote *Z. tritici* pathogenicity (Karki et al., 2021).

In wheat, twenty-four *Z. tritici* resistance genes (R genes) have been reported (Saintenac et al., 2021; Yang et al., 2022). *Stb6* encodes a conserved, wall-associated kinase (WAK) with an extracellular domain that recognizes the apoplastic AvrStb6 effector from *Z. tritici* (Brown et al., 2015; Saintenac et al., 2018; Zhong et al., 2017). AvrStb6 accumulates in epiphytic hyphae of *Z. tritici* in close proximity to the stomata, and its recognition triggers the induction of genes re-sponsible for stomatal closure in resistant cultivars (Alassimone et al., 2024). The resistance protein *Stb16q* also encodes a cysteine-rich, wall-associated-like receptor kinase that presumably recognizes a fungal carbohydrate to trigger a resistance response (Saintenac et al., 2021). Intriguingly, *Stb16q* confers full broad-spectrum resistance against 64 geographically diverse isolates of *Z. tritici* (Saintenac et al., 2021). Moreover, the *Stb9* resistance protein recognizes AvrStb9, a secreted, protease-like protein with a predicted S41 conserved domain and triggers an immune response following the gene-for-gene model (Amezrou et al., 2023). However, the function of *Stb9* is still not clear as it might be acting as a classical receptor protein, an inhibitor of AvrStb9 protease activity, or as the guard protein of the AvrStb9 target protein (Amezrou et al., 2023).

Recently, a genome-wide association study (GWAS) identified a 99-kb region containing six candidate genes and led to eventual cloning of the *Stb15* resistance gene. *Stb15* encodes an intronless G-type lectin receptor-like kinase (LecRK) with an intracellular serine/threonine receptor-like protein kinase and three extracellular domains (Hafeez et al., 2025). The G-type lectin domain of *Stb15* likely binds mannose and may play a role in basal plant immunity, as demonstrated previously in *Arabidopsis thaliana,* where lectins play a role in stomatal innate immunity responses (Singh and Zimmerli, 2013). *Stb15* could function as part of a guard/guardee pair, triggering isolate-specific resistance, or it may bind glycoproteins, similar to other LecRKs that interact with secreted proteins. A candidate for the corresponding *AvrStb15* gene, potentially a small, secreted protein, has been proposed, but further research is needed to confirm this interaction and its underlying mechanisms (Hafeez et al., 2025).

Non-host resistance (NHR) is commonly broad spectrum and, hence, likely more durable than that provided by classical R genes (Ayliffe & Sørensen, 2019; Gill et al., 2015). Several types of NHR involve a hypersensitive response, as is seen in qualitative resistance conferred by Avr recognition (Panstruga & Moscou, 2020). Evidence that Avr recognition is involved in NHR was found by studying the closest homologs of the *Z. tritici* Avr3D1 gene in sister fungal species that are nonpathogenic in wheat (Meile et al., 2023). The homologous genes induce a strong defense response in wheat when expressed in a virulent mutant strain of *Z. tritici*, suggesting they contribute to establishing host range and are involved in NHR (Meile et al., 2023). Barley (*Hordeum vulgare* subsp. *vulgare*) is closely related to wheat and is infected by *Z. passerinii* (Feurtey et al., 2020; Goodwin & Zismann, 2001), a species that is closely related to *Z. tritici.* Barley is a non-host of *Z. tritici* and thus shows strong resistance to this pathogen. Identifying and comparing DEGs in *Z. tritici* during a non-host interaction with barley is central to testing whether similar virulence factors and molecular pathways are expressed during susceptible, resistant (R gene) and non-host interactions. In addition, studying pathogen gene expression during a non-host interaction with a closely related grass species of wheat can identify candidate effectors that determine host specificity, and may lead to the identification of host target proteins. This information can be transferred from barley into the *Z. tritici* – wheat pathosystem to provide additional sources for breeding resistance in wheat.

In the present study, we investigate pathogen gene responses during interactions with susceptible, resistant, and non-host cultivars, using a time course of transcriptomic data covering distinct phases of the infection process. We examine three types of interaction: susceptible interaction with wheat cultivar Taichung 29; two resistant interactions with wheat cultivars Veranopolis and Israel 493; and a non-host interaction with barley to test the hypothesis that *Z. tritici* expresses different sets of genes tailored to each type of interaction. We identified thirty-one putative effectors that are upregulated during early disease development in a susceptible interaction compared to a non-host interaction. From these, six candidate effectors were selected based on their expression profiles across time points in the replicates of the susceptible interaction, and their subcellular localization was investigated using *Agrobacterium*-mediated transient expression in *Nicotiana benthamiana*.

## Results

### Gene expression analysis during the interaction of *Z. tritici* and its wheat host

Differential gene expression in *Z. tritici* was determined between three types of host interactions: (1) a susceptible interaction with the wheat cultivar Taichung 29; (2) a resistant interaction with the wheat cultivar Veranopolis, which carries the *Stb2* and *Stb6* resistance genes; and (3) a resistant interaction with the wheat cultivar Israel 493, which carries the *Stb3* and *Stb6* resistance genes (Table 1). Comparison of pathogen gene expression in the susceptible host versus the resistant hosts, allowed us to identify genes that are necessary for the transition from biotrophic to necrotrophic growth. We measured the gene expression at six time points: 1, 3, 6, 10, 17 and 23 days post inoculation (DPI). Previous studies have established that the transition from biotrophic to necrotrophic growth in *Z. tritici* takes place at approximately 10 DPI (Palma-Guerrero et al., 2017; Rudd et al., 2015; Yang et al., 2013).

**Table 1.** Comparisons of gene expression by the wheat pathogen *Zymoseptoria tritici* during susceptible and resistant interactions on wheat and its non-host interaction on barley.

| Host inoculated | Compared to | Type of comparison |
| --- | --- | --- |
| Taichung 29 wheat | Veranopolis (*Stb2*) wheat | Susceptible vs. resistant |
|  | Israel 493 (*Stb3*) wheat | Susceptible vs. resistant |
|  | Kindred barley | Susceptible vs. Non host |
| Veranopolis (*Stb2*) wheat | Israel 493 (*Stb3*) wheat | Resistant vs. resistant |
| Kindred barley | Veranopolis (*Stb2*) wheat | Non host vs. resistant |
|  | Israel 493 (*Stb3*) wheat | Non host vs. resistant |

Close to two billion reads were collected for the susceptible interaction and resistant interaction with cultivar Veranopolis, in total, over all time points and replications, while more than two billion reads were obtained for the resistant interaction with cultivar Israel 493 (Supplementary Table 1). At 1 and 3 DPI, the percentage of reads mapped to the fungal genome relative to the total was similar between the susceptible and resistant interactions. However, in the susceptible interaction, this percentage increased at 6 and 10 DPI, aligning with the expected rise in fungal biomass preceding the transition to necrotrophy. In contrast, for both resistant interactions, at 6 and 10 DPI, the proportion of fungal reads remained comparable to the levels observed at 3 DPI (Supplementary Table 1). At 17 and 23 DPI, the susceptible interaction with cultivar Taichung 29 showed a significant increase in fungal reads, corresponding to the proliferation of macroscopic symptoms on leaves. Conversely, in both resistant interactions, the numbers of reads mapped to the *Z. tritici* genome remained at the lowest levels, consistent with the absence of visible symptoms and the limited proliferation of fungal biomass in the resistant cultivars (Adhikari et al., 2025). Overall, at 1, 3 and 6 DPI, only a small percentage (less than 1%) of the total reads aligned to the fungal genome in both susceptible and resistant interactions. However, from 10 DPI onward, the proportion of mapped fungal reads increased in the susceptible interaction, reaching its peak at 23 DPI, where it exceeded 40% in two replicates.

Gene expression patterns in the susceptible and the two resistant interactions were explored with principal component analysis (PCA). The PCA plots for 1, 3, 6 and 10 DPI (Fig. 1) and for 17 and 23 DPI (Fig. 2.) were generated separately because normalization had to be performed in two groups due to the very few reads in the resistant interactions at 17 and 23 DPI (see Materials and Methods). PCA shows that gene expression is very similar in the susceptible and resistant interactions early during infection (1 and 3 DPI) (Fig. 1.). Most replicate samples are well clustered, indicating limited biological variation within replicate time points. At 6 DPI, gene expression in the susceptible and resistant interactions begins to diverge, and at 10 DPI, the susceptible interaction is distinctly separated from the resistant interactions, primarily along principal component 2, while gene expression in the samples from both resistant interactions at 10 DPI remains similar to that at 6 DPI.

**Fig. 1.**
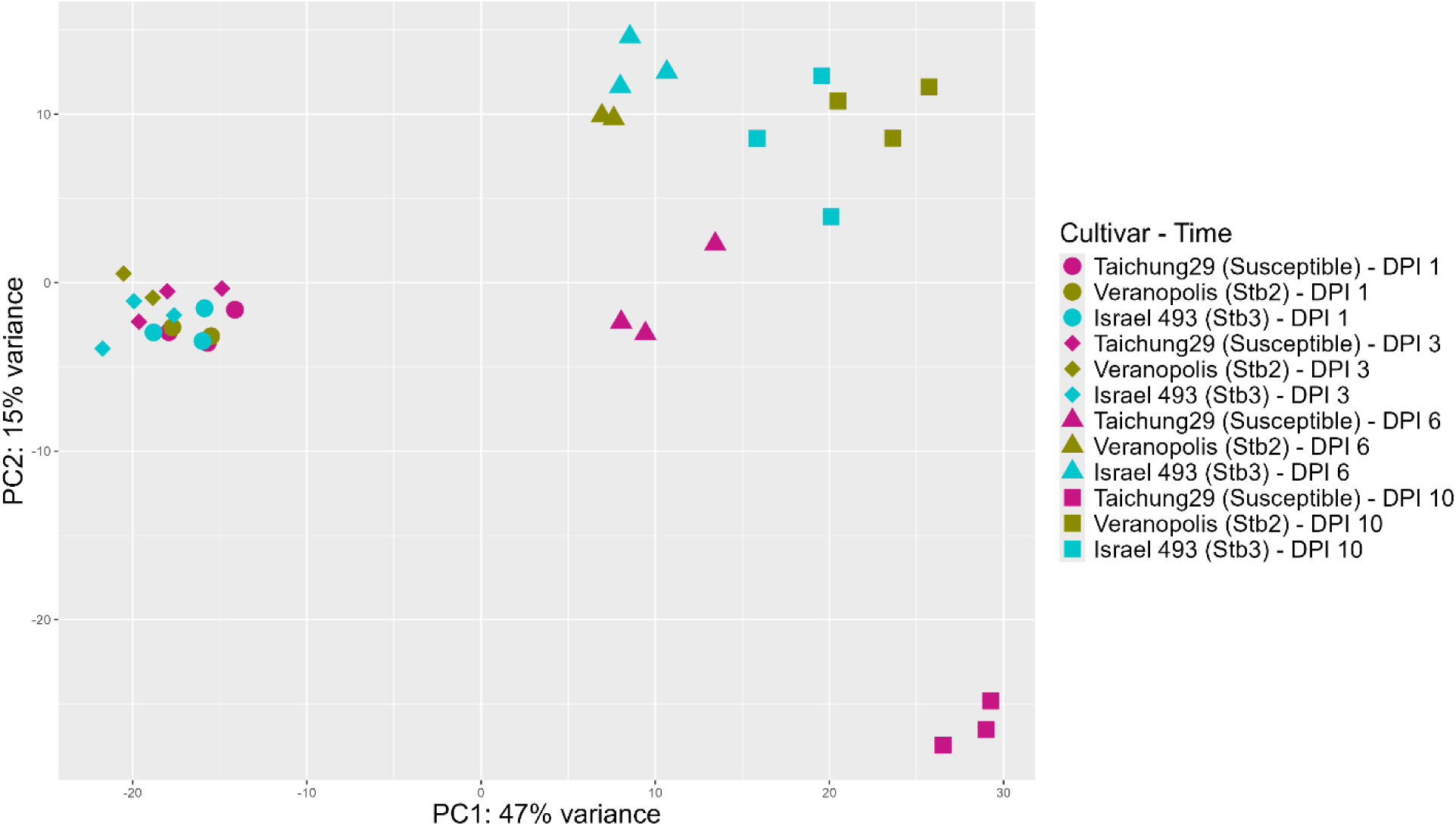
Principal component analysis (PCA) of the normalized and filtered counts for RNA sequencing reads of *Zymoseptoria tritici* inoculated onto the highly susceptible wheat cultivar Taichung 29 (Susceptible) or the R-gene-containing resistant wheat cultivars Veranopolis (*Stb2* resistance gene) or Israel 493 (*Stb3*) at 1, 3, 6, and 10 days post inoculation (DPI).

**Fig. 2.**
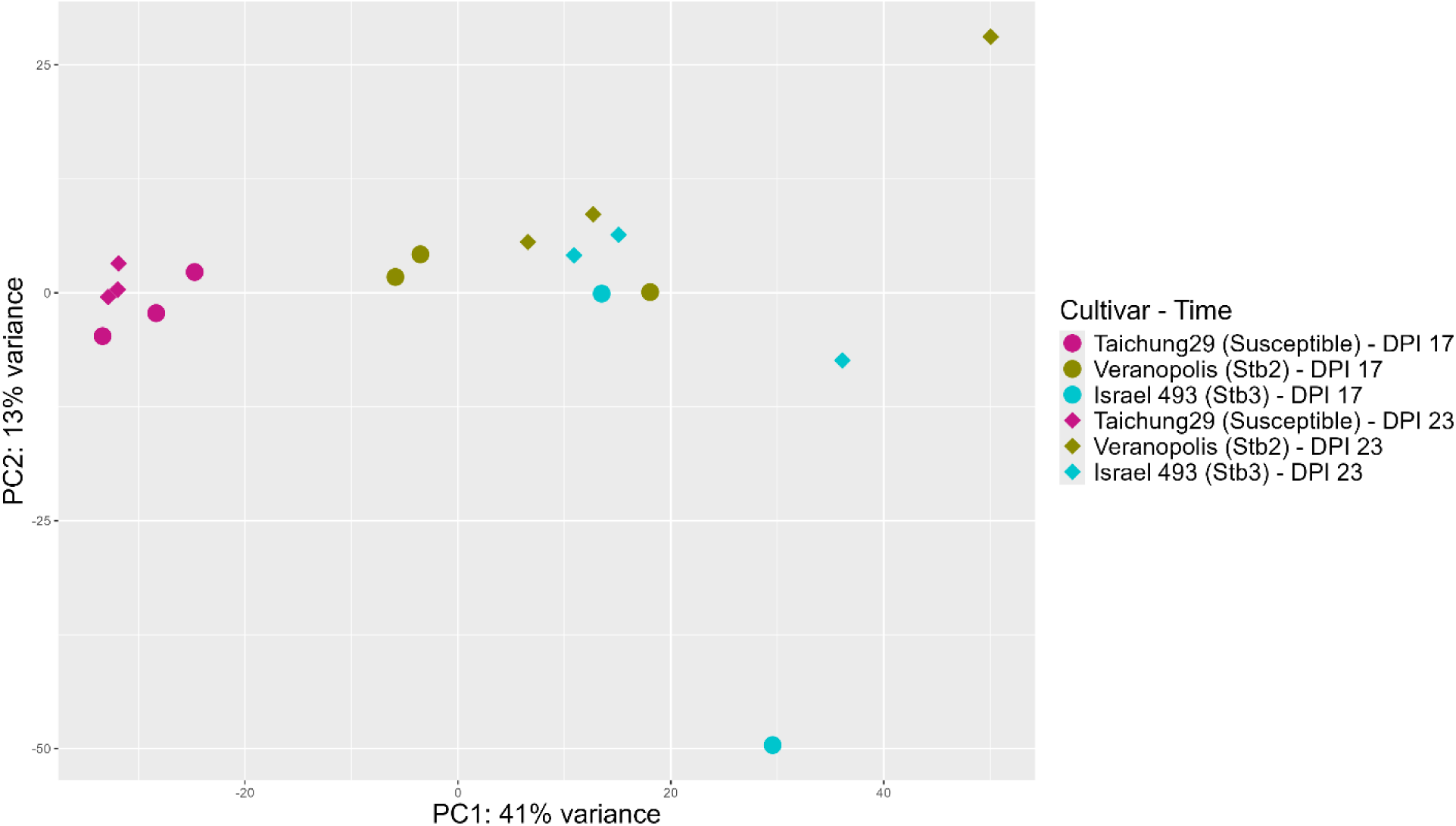
Principal component analysis (PCA) of the normalized and filtered counts for RNA sequencing reads of *Zymoseptoria tritici* inoculated onto the highly susceptible wheat cultivar Taichung 29 (Susceptible) or the R-gene-containing resistant wheat cultivars Veranopolis (*Stb2* resistance gene) or Israel 493 (*Stb3*) at 17 and 23 days post inoculation (DPI).

A similar pattern is observed at 17 and 23 DPI (Fig. 2), where the susceptible interaction forms a separate cluster from the resistant interactions. In both susceptible and resistant interactions, the 17 and 23 DPI points cluster together for each wheat variety, suggesting that there is little change in gene expression from 17 to 23 DPI, although the susceptible and resistant interactions remain clearly distinguished. The replicate points for the resistant interactions show considerable variability due to the low number of read counts obtained at late infection stages in the resistant interactions, but the trend is clear.

The highest number of DEGs in *Z. tritici* was observed at 10 DPI when comparing the susceptible interaction on the cultivar Taichung 29 with both resistant interactions on the *Stb2*-carrying cultivar Veranopolis (601 DEGs) and on the *Stb3-*carrying cultivar Israel 493 (587 DEGs) (Table 2). A total of 275 and 226 genes in *Z. tritici* was found to be up-regulated at 10 DPI during the susceptible interaction compared to the resistant *Stb2* and *Stb3* interactions, respectively. The rapid and pronounced activation of a large number of DEGs in *Z. tritici* at 10 DPI in the susceptible interaction corresponds with the first visible symptoms on the leaves of the susceptible cultivar Taichung 29 (Supplementary Fig. S1) and marks the transition from the biotrophic to the necrotrophic stage. Interestingly, a total of 326 and 361 genes in *Z. tritici* were identified as down-regulated in the susceptible interaction at 10 DPI when compared to the resistant interactions with Veranopolis and Israel 493, respectively. Since the resistant interactions were used as the baseline for these comparisons, this means that these same genes were up-regulated in the resistant interactions relative to the susceptible interaction. This suggests that during the resistant interactions, *Z. tritici* activates a larger set of genes, potentially as part of a counteractive response to the defense mechanisms deployed by the resistant cultivars at 10 DPI.

**Table 2.** Differentially expressed genes (DEGs) of the wheat pathogen *Zymoseptoria tritici* when comparing three types of host interaction^a^.

| Comparison | Time point<sup>b</sup> | Up<sup>c</sup> | Down<sup>c</sup> | Total DEGs | Total detectable genes |
| --- | --- | --- | --- | --- | --- |
| **Resistant (Stb2) – Resistant (Stb3)<sup>d</sup>** | 1 | 0 | 0 | 0 | 1,504 |
|  | 3 | 0 | 0 | 0 |  |
|  | 6 | 1 | 0 | 1 |  |
|  | 10 | 1 | 11 | 12 |  |
|  | 17 | 13 | 1 | 14 | 222 |
|  | 23 | 3 | 0 | 3 |  |
| **Susceptible - Resistant (Stb2)<sup>e</sup>** | 1 | 0 | 0 | 0 | 1,504 |
|  | 3 | 1 | 4 | 5 |  |
|  | 6 | 12 | 0 | 12 |  |
|  | 10 | 275 | 326 | 601 |  |
|  | 17 | 21 | 11 | 32 | 222 |
|  | 23 | 33 | 23 | 56 |  |
| **Susceptible - Resistant (Stb3)<sup>f</sup>** | 1 | 1 | 1 | 2 | 1,504 |
|  | 3 | 0 | 4 | 4 |  |
|  | 6 | 21 | 0 | 21 |  |
|  | 10 | 226 | 361 | 587 |  |
|  | 17 | 34 | 13 | 47 | 222 |
|  | 23 | 34 | 22 | 56 |  |

<sup>a</sup> DEGs were identified for the first-listed condition in each comparison relative to the second-listed condition, which served as the baseline. Horizontal lines within each comparison indicate normalization groups. Gene counts for time points 1, 3, 6, and 10 DPI were normalized together, while counts for time points 17 and 23 DPI were normalized separately as a distinct group.
<sup>b</sup> 6-point time course. Time points are measured in days after inoculation.
<sup>c</sup> Significantly differentially expressed genes that are up- or down-regulated at a False Discovery Rate (FDR) of 5%.
<sup>d</sup> Responses of *Z. tritici* isolate IPO323 when grown on *Stb2*-resistant plants of cultivar Veranopolis compared to those on *Stb3*-resistant plants of cultivar Israel 493
<sup>e</sup> Responses of *Z. tritici* isolate IPO323 when grown on susceptible cultivar Taichung 29 compared to those on *Stb2*-resistant plants of cultivar Veranopolis
<sup>f</sup> Responses of *Z. tritici* isolate IPO323 when grown on susceptible cultivar Taichung 29 compared to those on *Stb3*-resistant plants of cultivar Israel 493

We determined the number of up-regulated genes in *Z. tritici* that were in common among the comparisons between the susceptible interaction and both resistant interactions. The Venn diagram (Fig. 3A) represents the two sets of up-regulated genes. The intersection of these two sets includes 183 (57.5%) genes that are up-regulated in both comparisons. These genes are likely part of core mechanisms of interaction with a susceptible host at 10 DPI, independent of which resistant interaction is used as the control. This suggests that these genes are likely involved in the common molecular mechanisms or pathways in the pathogen that are triggered during the susceptible interaction at 10 DPI. In addition to their potential role in mediating the transition to the necrotrophic stage of infection, the fungal genes that are upregulated in susceptible compared to both resistant interactions may also represent targets of host-mediated suppression. Their reduced expression in the interaction with resistant cultivars provides evidence of the direct and indirect effects of the presence of the resistance genes in those cultivars. Similarly, in the Venn diagram (Fig. 3B), the intersection includes 234 genes that are down-regulated (51.7%) in the susceptible interaction compared to both resistant interactions at 10 DPI. Since the resistant interactions serve as the baseline, these same genes are up-regulated in both resistant interactions relative to the susceptible interaction. This shared gene set likely represents a core transcriptional response in *Z. tritici* to counteract the defense mechanisms activated by resistant cultivars, suggesting a common response against a resistant cultivar regardless of the specific composition of the resistant genotype.

**Fig. 3.**
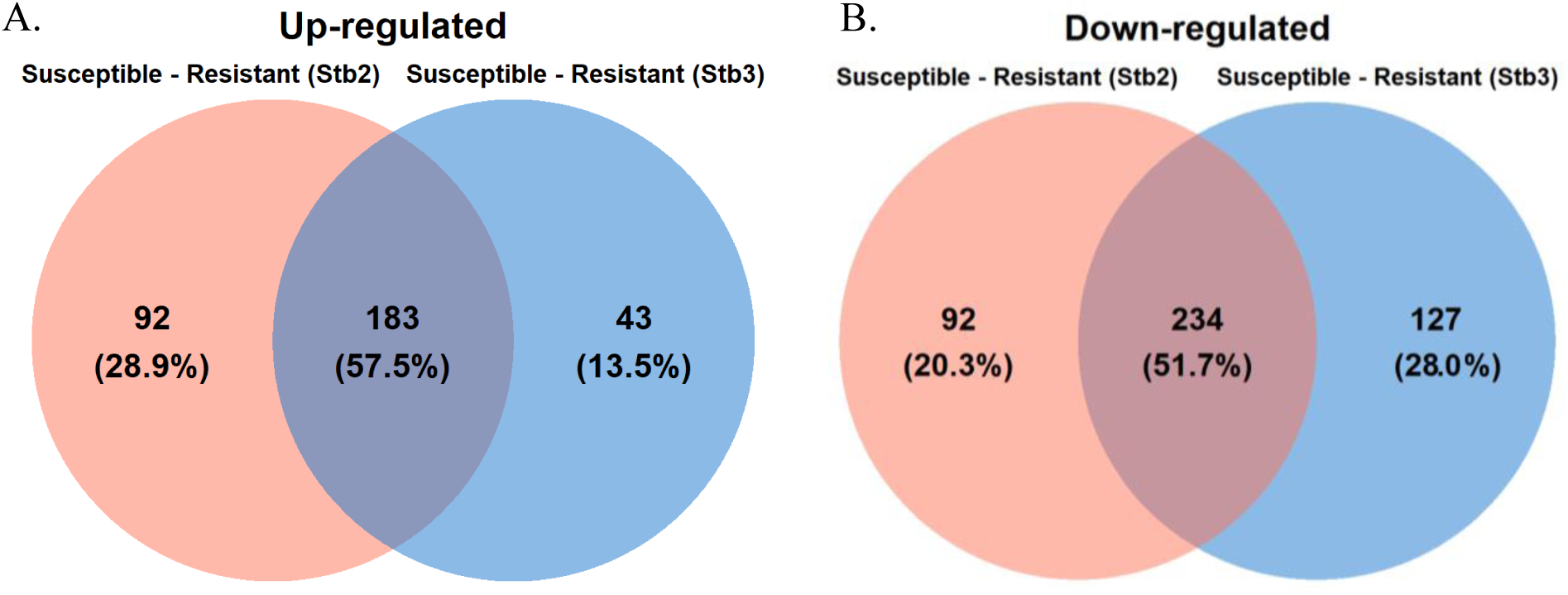
Venn diagrams showing common and unique differentially expressed genes of *Zymoseptoria tritici* at 10 days post inoculation (DPI) in the susceptible interaction compared to the two resistant interactions on wheat. (A) Up-regulated genes shared between the comparisons of the susceptible interaction versus the resistant interactions with cultivars Veranopolis (*Stb2*) and Israel 493 (*Stb3*). The intersection represents 183 (57.5%) genes that are up-regulated in the pathogen during the susceptible interaction relative to both resistant interactions. (B) Down-regulated genes shared between the comparisons of the susceptible interaction versus the resistant interactions with cultivars Veranopolis (*Stb2*) and Israel 493 (*Stb3*). The intersection represents 234 (51.7%) genes that are down-regulated in the pathogen during the susceptible interaction relative to both resistant interactions.

We identified genes that were exclusively expressed at certain time points in the susceptible interaction compared to the resistant interactions on cultivars Veranopolis and Israel 493. Remarkably, 249 up-regulated DEGs were uniquely expressed at 10 DPI in the susceptible interaction compared to the resistant interaction with Veranopolis (Fig. 4A). The same pattern of uniquely differentially expressed genes was observed at 10 DPI in the comparison of *Z. tritici* grown on the susceptible cultivar Taichung 29 compared to the resistant cultivar Israel 493 (Fig. 4B). These results demonstrate that the number of DEGs that are exclusively activated in *Z. tritici* at 10 DPI is remarkably high compared to other time points, which indicates that a defined reprogramming in gene expression occurs at this stage.

**Fig. 4.**
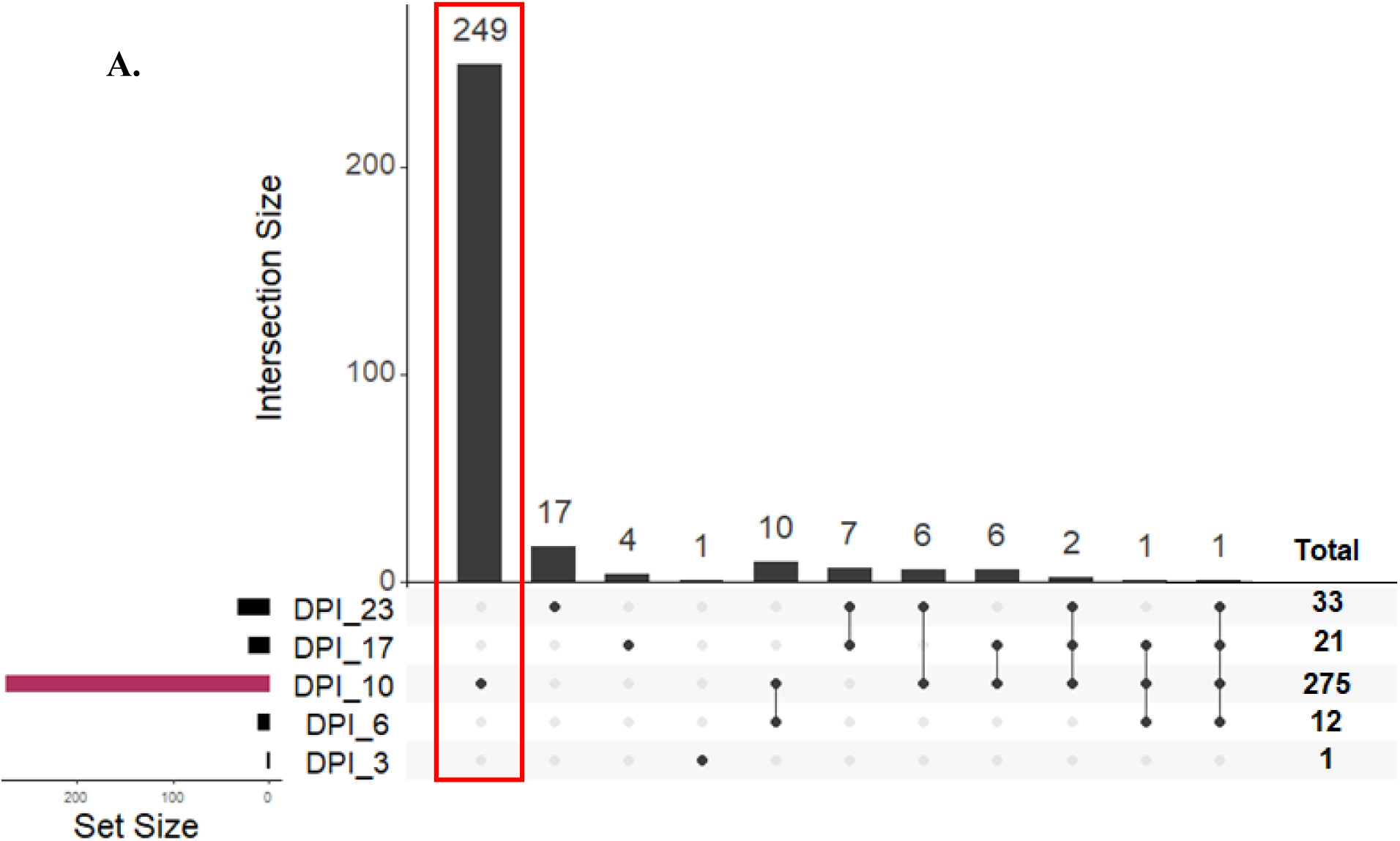
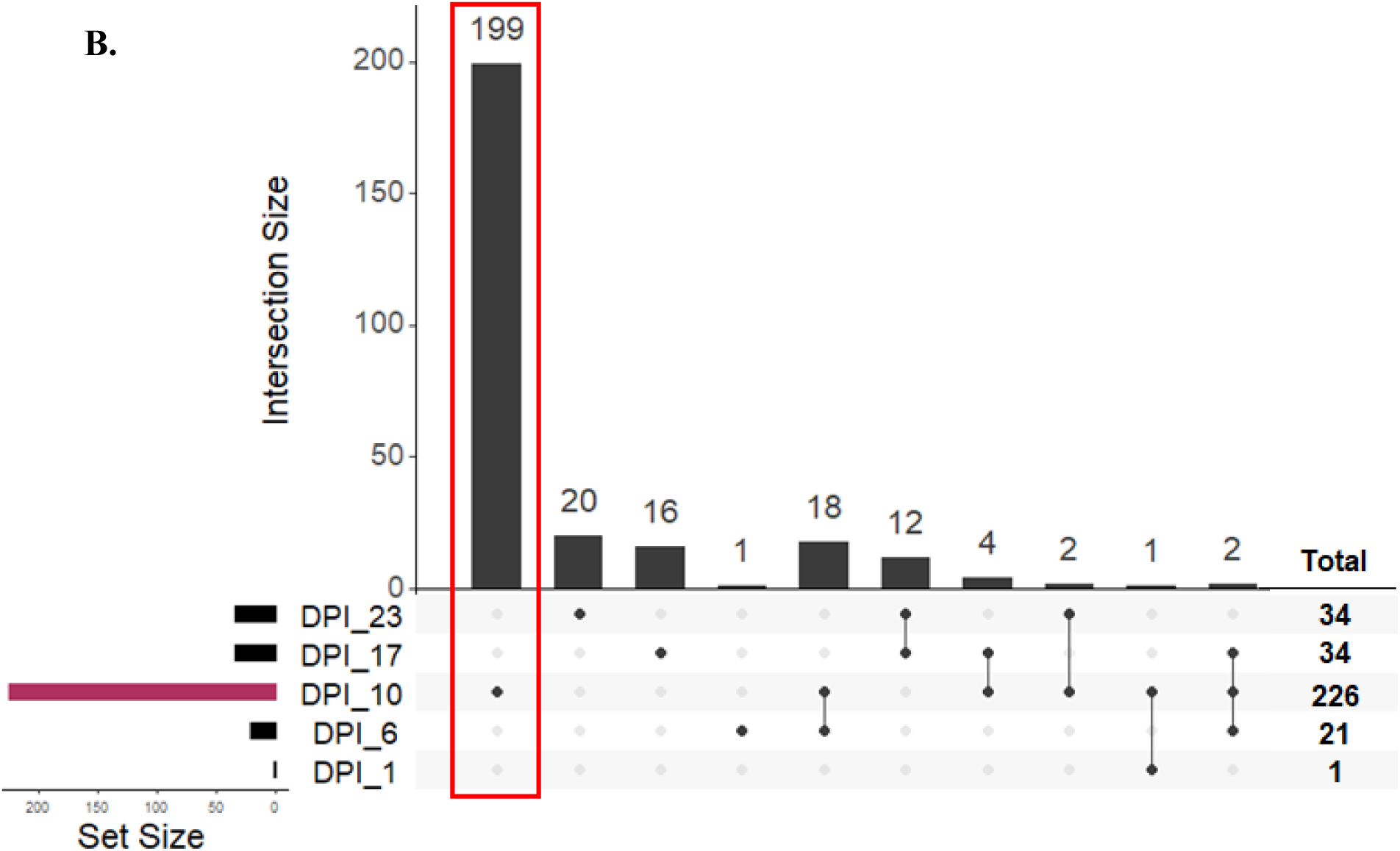
Numbers of unique and shared up-regulated differentially expressed genes (DEGs) of the wheat pathogen *Zymoseptoria tritici* during the susceptible interaction on the cultivar Taichung 29 compared to the resistant interaction on: A) the *Stb2*-carrying cultivar Veranopolis; and to the resistant interaction on B) the *Stb3-*carrying cultivar Israel 493; at 1, 3, 6, 10, 17 and 23 days post inoculation (DPI). The colored horizontal bar indicates the total number of up-regulated DEGs at the time point showing the highest number of uniquely up-regulated genes. The red rectangle highlights the number of genes that were exclusively differentially up-regulated at 10 DPI.

To identify differences in pathogen gene expression in the two resistant interactions, we determined DEGs in *Z. tritici* during the resistant interaction with Veranopolis compared to the resistant interaction with Israel 493. Only 1 DEG was up-regulated in *Z. tritici* inoculated on the Veranopolis cultivar compared to Israel 493 at 6 DPI. In total 12, 14 and 3 genes were found to be differentially expressed at 10, 17 and 23 DPI (Table 2), respectively. No DEGs were identified when comparing the two resistant interactions at 1 and 3 DPI. This shows that *Z. tritici* follows a nearly identical pattern of gene expression during its interaction with both resistant wheat cultivars.

### Gene Ontology and enrichment analysis during host interactions

*Zymoseptoria tritici* DEGs were grouped according to their putative roles in biological processes (BP), molecular functions (MF) and cellular components (CC) as established by the Gene Ontology Consortium (http://www.geneontology.org/). The enriched GO terms in the up-regulated genes in *Z. tritici* at 10 DPI (Fig. 5) collectively indicate that the pathogen is actively metabolizing and synthesizing key macromolecules, such as proteins, nucleic acids, and nitrogenous compounds. This is suggested by the enrichment of GO terms associated with macromolecular metabolic and biosynthetic processes at 10 DPI (Fig. 5) including organonitrogen compound metabolic process (GO:1901564), ribonucleoprotein complex biogenesis (GO:0022613), and pyrimidine-containing compound biosynthetic process (GO:0072528). Numerous GO terms related to ribosomal components are enriched at 10 DPI in the susceptible interaction compared to the resistant interactions, suggesting that the pathogen is actively ramping up protein synthesis. This is likely to support the pathogen’s metabolic needs as it establishes a necrotrophic infection. The enrichment of GO terms in the categories nucleic acid binding (GO:0003676) and rRNA binding (GO:0019843) points to upregulation of processes related to RNA processing and stability, which may further support the pathogen’s increased need for protein synthesis and efficient gene expression. At 17 DPI, the enrichment of GO categories related to transport activities (Fig. 5), suggests that the pathogen is likely focusing on regulating the ion homeostasis and controlling the cellular uptake and efflux of ions. These processes are likely linked to nutrient acquisition, as ion transporters are essential for maintaining osmotic balance, acquiring nutrients, and detoxifying the pathogen’s environment (Barata-Antunes et al., 2021). Such activities might be crucial as the pathogen progresses into later stages of infection, facilitating its adaptation and survival. It might also suggest that *Z. tritici* is switching from internal synthesis to importing metabolites from the breakdown of host cells.

**Fig. 5.**
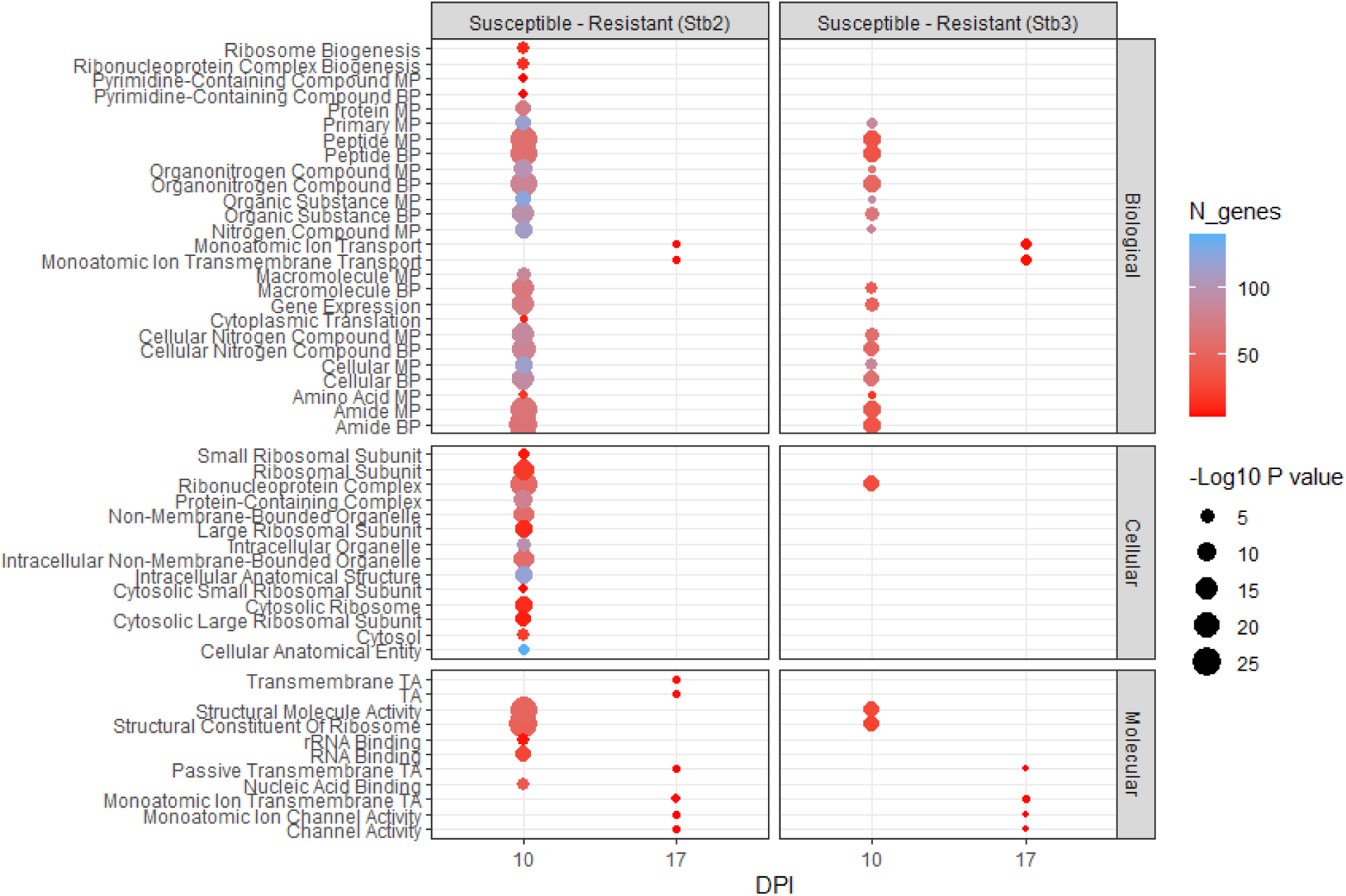
Gene Ontology (GO) terms enriched at 10 and 17 days post inoculation (DPI) in the susceptible interaction compared to the resistant interactions on the *Stb2-*contining cultivar Veranopolis (left panel) and on the *Stb3-*containing cultivar Israel 493 (right panel) following inoculation with the wheat pathogen *Zymoseptoria tritici*. Only up-regulated genes were included in the GO analysis. The number of genes included in the GO analysis is 207 for the susceptible interaction compared to the resistant interaction on the *Stb2*-containing cultivar Veranopolis at 10 and 17 DPI, and 166 for the susceptible interaction compared to the resistant interaction on the *Stb3*-containing cultivar Israel 493 at 10 and 17 DPI. Metabolic process (MP), Biosynthetic process (BP), Transporter Activity (TA).

The KEGG pathway enrichment analysis showed that 13 pathways were significantly enriched in the up-regulated genes at 10 DPI during the susceptible interaction compared to both resistant interactions. The top five enriched pathways were carbon metabolism, biosynthesis of antibiotics, biosynthesis of secondary metabolites, glycolysis and gluconeogenesis. In contrast, at 23 DPI only 3 pathways were enriched among the up-regulated genes in the susceptible interaction compared to the resistant interactions including carbon metabolism, biosynthesis of antibiotics, and glycolysis and gluconeogenesis.

In the susceptible interaction compared to the resistant interactions, we observed a significant up-regulation of genes at 10 DPI that encode carbohydrate-active enzymes (CAZymes). Among these, we identified eight glycoside hydrolases (GHs) and one carbohydrate esterase (CE), all of which are cell wall-degrading enzymes (CWDEs). Additionally, our analysis revealed 10 glycosyltransferases (GTs) and 10 enzymes with auxiliary activity (AA) among the up-regulated genes (Supplementary Fig. S2). Families of AA enzymes involved in oxidation of aromatic compounds and fungal growth such as the multicopper oxidase family (AA1), and the alcohol oxidase subfamily (AA3_3) were up-regulated at 10 DPI in both comparisons between the susceptible interaction and the two resistant interactions.

Interestingly, among the down-regulated genes at 10 DPI in the susceptible interaction using the resistant interactions as baseline, we also identified 12 GHs, 8 GTs, and 11 enzymes in the AA category. Families of AA enzymes involved in detoxification, oxidation of compounds, and fungal growth such as the benzoquinone reductase family (AA6), the alcohol dehydrogenase subfamily (AA3_2) and the alcohol oxidase subfamily (AA3_3) were down-regulated in the susceptible interaction compared to both resistant interactions. This means that these families of enzymes were up-regulated in both resistant interactions. This observation indicates the likely production of CAZymes in the pathogen at the transition stage during a resistant interaction, possibly to counteract resistant responses in the wheat cultivars. For instance, the CWDEs secreted by different plant pathogens can act as effectors, and they can inhibit or elicit plant immunity (Gu et al., 2024; Gui et al., 2017; Zhang et al., 2021).

We identified predicted small, secreted proteins (SSPs) among DEGs at 10 DPI. Notably, 26 SSPs were down-regulated in the susceptible interaction compared to both resistant interactions. Additionally, 22 SSPs were up-regulated in the susceptible interaction relative to the resistant interaction on the cultivar Veranopolis, and 23 SSPs showed up-regulation in the susceptible interaction compared to the resistant interaction on the cultivar Israel 493.

### Gene expression during a non-host interaction

Differential gene expression in *Z. tritici* was determined in three host interactions compared to a non-host interaction with the barley cultivar Kindred (Table 1). Comparison of pathogen gene expression in the susceptible and resistant hosts, versus the non-host species, allows us to identify genes involved in initial colonization and host recognition.

The percentage of reads mapped to the fungal genome during the non-host interaction remained consistently low across all time points, showing no significant increase over time. It never exceeded 0.2% and, in most replicates, ranged between 0.02% and 0.06% (Table 2). As a result, only a limited set of genes could be detected in comparisons using the non-host interaction as the baseline treatment.

In contrast to the comparison between the susceptible interaction and the resistant interactions, the largest differences between the susceptible and the non-host interaction are seen at 3 DPI. There were 171 DEGs in *Z. tritici,* of which 124 were up-regulated and 47 were down-regulated (Table 3) in the susceptible interaction compared to the non-host interaction with *H. vulgare* used as the baseline. The genes that are up-regulated in the susceptible interaction compared to the non-host interaction may play a role in initially colonizing the susceptible host. There were 154 DEGs in the resistant interaction between *Z. tritici* and the Veranopolis (*Stb2*) resistant cultivar compared to the non-host interaction (Table 3). This is similar to the 160 DEGs in *Z. tritici* in the resistant interaction between *Z. tritici* and the Israel 493 (*Stb3*) cultivar compared to the non-host interaction used as the baseline (Table 3).

**Table 3.** Differentially expressed genes (DEGs) of the wheat pathogen *Zymoseptoria tritici* when comparing susceptible and resistant interactions versus a non-host interaction on barley^a^.

| Comparison | Time point<sup>b</sup> | Up<sup>c</sup> | Down<sup>c</sup> | Total DEGs | Total detectable genes |
| --- | --- | --- | --- | --- | --- |
| **Susceptible–non host<sup>d</sup>** | 1 | 74 | 40 | 114 | 284 |
|  | 3 | 123 | 47 | 171 |  |
|  | 6 | 43 | 48 | 91 | 352 |
|  | 10 | 94 | 52 | 146 |  |
|  | 17 | 57 | 14 | 71 | 107 |
|  | 23 | 63 | 17 | 80 |  |
| **Resistant (*Stb2*) – non host<sup>e</sup>** | 1 | 67 | 40 | 107 | 284 |
|  | 3 | 107 | 47 | 154 |  |
|  | 6 | 40 | 44 | 84 | 352 |
|  | 10 | 68 | 41 | 109 |  |
|  | 17 | 20 | 9 | 29 | 107 |
|  | 23 | 30 | 27 | 57 |  |
| **Resistant (*Stb3*) – non host<sup>f</sup>** | 1 | 68 | 41 | 109 | 284 |
|  | 3 | 108 | 52 | 160 |  |
|  | 6 | 54 | 59 | 113 | 352 |
|  | 10 | 70 | 39 | 109 |  |
|  | 17 | 9 | 9 | 18 | 107 |
|  | 23 | 22 | 23 | 45 |  |

<sup>a</sup> DEGs were identified for the first-listed condition in each comparison relative to the second-listed condition, which served as the baseline. Horizontal lines within each comparison indicate normalization groups. Gene counts for time points 1 and 3 DPI were normalized together, 6 and 10 DPI were normalized as a separate group, and 17 and 23 DPI were normalized together as another distinct group.
<sup>b</sup> 6-point time course. Time points are measured days after inoculation.
<sup>c</sup> Significantly differential expressed genes that are up- or down-regulated at a False Discovery Rate (FDR) of 5%.
<sup>d</sup> Responses of *Z. tritici* isolate IPO323 when grown on plants of susceptible wheat cultivar Taichung 29 compared to those on non-host barley plants
<sup>e</sup> Responses of *Z. tritici* isolate IPO323 when grown on *Stb2*-resistant plants of cultivar Veranopolis compared to those on non-host barley plants
<sup>f</sup> Responses of *Z. tritici* isolate IPO323 when grown on *Stb3*-resistant plants of cultivar Israel 493 compared to those on non-host barley plants

The numbers of up-regulated DEGs at 10 DPI show a slight increase in contrast to the previous time point at 6 DPI when comparing both the susceptible and resistant interactions against the non-host interaction. At 17 and 23 DPI, for the genes for which we were able to retrieve measurable counts, we observed higher numbers of DEGs in the susceptible interaction than the numbers of DEGs at 17 and 23 DPI in the resistant interactions compared to the nonhost interaction. The fact that we were able to identify very little fungal RNA originating from the *Z. tritici* genome at 17 and 23 DPI, in both resistant and non-host interactions, also reveals that there is a significant reduction in the fungal biomass compared to 1 to 10 DPI in these interactions and compared to the susceptible interaction. This also could mean that the fungus is not transcriptionally active at 17 and 23 DPI in resistant and non-host interactions.

We determined the number of up-regulated genes in *Z. tritici* that were in common in the three comparisons between the susceptible and the two resistant host interactions compared to the non-host interaction. The Venn diagram (Fig. 6A) illustrates the three sets of up-regulated genes, with an intersection of 98 genes (74.8%) that are consistently up-regulated in the pathogen in the comparisons of susceptible versus non-host, resistant (Veranopolis) versus non-host, and resistant (Israel 493) versus non-host. These genes likely have a role in core mechanisms involved in host interaction at 3 DPI, irrespective of whether the host is susceptible or resistant, i.e., genes that are important for the initial biotrophic growth phase. This suggests that these genes are part of fundamental host recognition pathways shared between both susceptible and resistant interactions at this early stage.

**Fig. 6.**
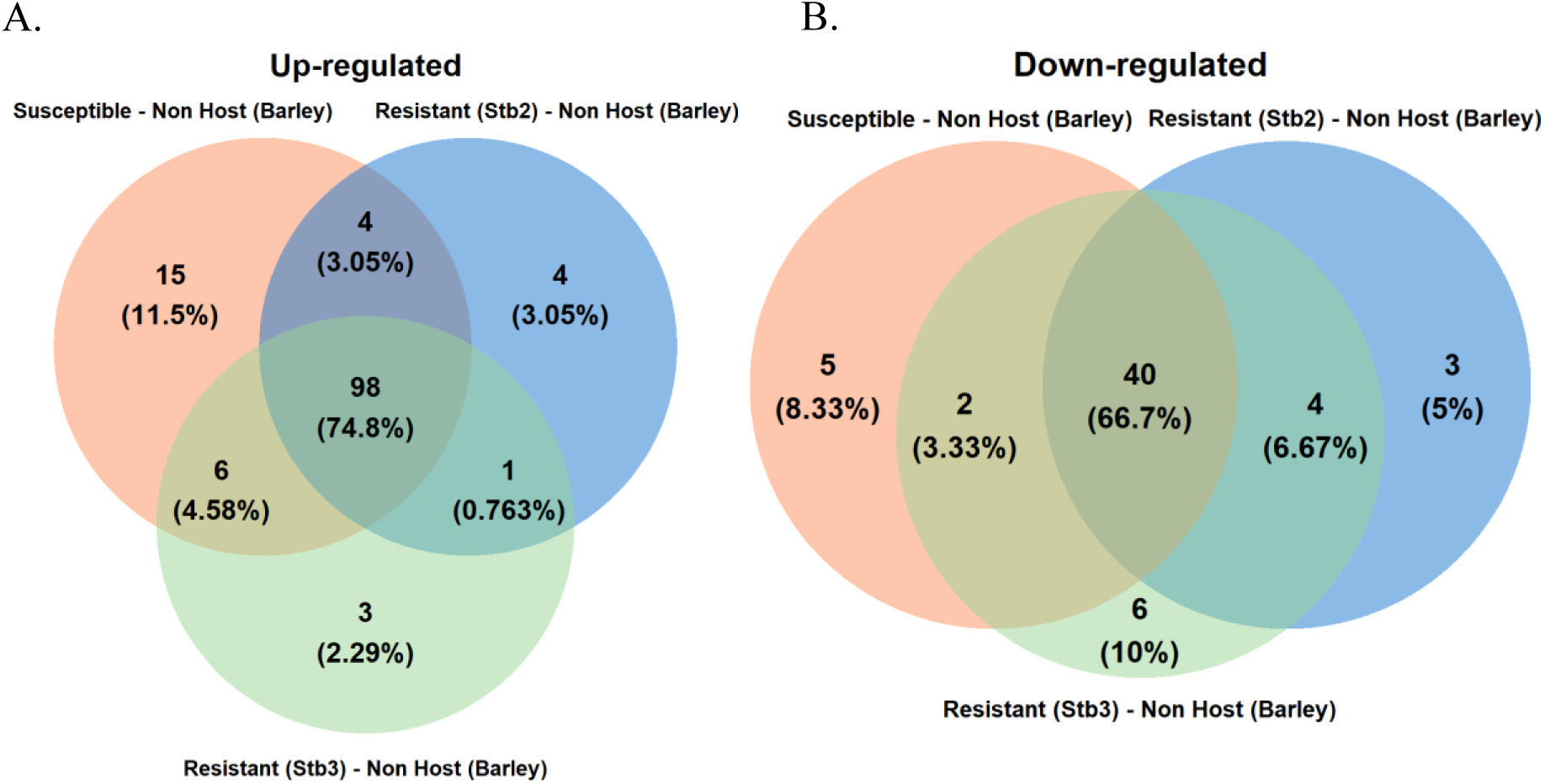
Venn diagrams showing common and unique differentially expressed genes of *Zymoseptoria tritici* at 3 days post inoculation (DPI) in the susceptible and resistant interactions on wheat compared to the non-host interaction on barley. (A) Up-regulated genes shared between the comparisons of the susceptible interaction versus the non-host interaction, the resistant interaction with Veranopolis (*Stb2*) versus the non-host interaction, and the resistant interaction with Israel 493 (*Stb3*) versus the non-host interaction. The intersection represents 98 (74.8%) genes that were up-regulated in the pathogen during the susceptible and resistant interactions relative to the non-host interaction. (B) Down-regulated genes shared between the comparisons of the susceptible interaction versus the non-host interaction, the resistant interaction with Veranopolis (*Stb2*) versus the non-host interaction, and the resistant interaction with Israel 493 (*Stb3*) versus the non-host interaction. The intersection represents 40 (66.7%) genes that were down-regulated in the pathogen during the susceptible and resistant interactions relative to the non-host interaction.

Similarly, for the down-regulated genes, the Venn diagram (Fig. 6B) shows an intersection of 40 genes (66.7%) shared across the three host interactions compared to the non-host interaction at 3 DPI. This indicates a common response to the interaction with barley, the non-host species, regardless of whether the control host interaction is susceptible or resistant. These common down-regulated genes reflect differences in the interaction and growth of *Z. tritici* on a non host at the early infection stages.

To find unique up-regulated DEGs that are activated at certain time points during the susceptible and resistant interactions compared to the non-host interaction and differentiate them from those that were expressed steadily, we determined unique and shared up-regulated DEGs between the sampling days. In the susceptible interaction compared to the non-host interaction, there were 52 up-regulated DEGs that were unique at 3 DPI, followed by 50 up-regulated DEGs shared at 1 and 3 DPI (Fig. 7A). There were 45 unique DEGs in *Z. tritici* at 3 DPI, followed by 45 DEGs at both 1 and 3 DPI when comparing the resistant interaction on Veranopolis (*Stb2*) with the non-host interaction (Fig. 7B). The combined number of up-regulated DEGs at 1 and 3 DPI is much higher than the unique genes at other time points, which indicates that the main difference in the pattern of gene activation between the resistant interaction and the non-host interaction occurs at the beginning of the infection process.

**Fig. 7.**
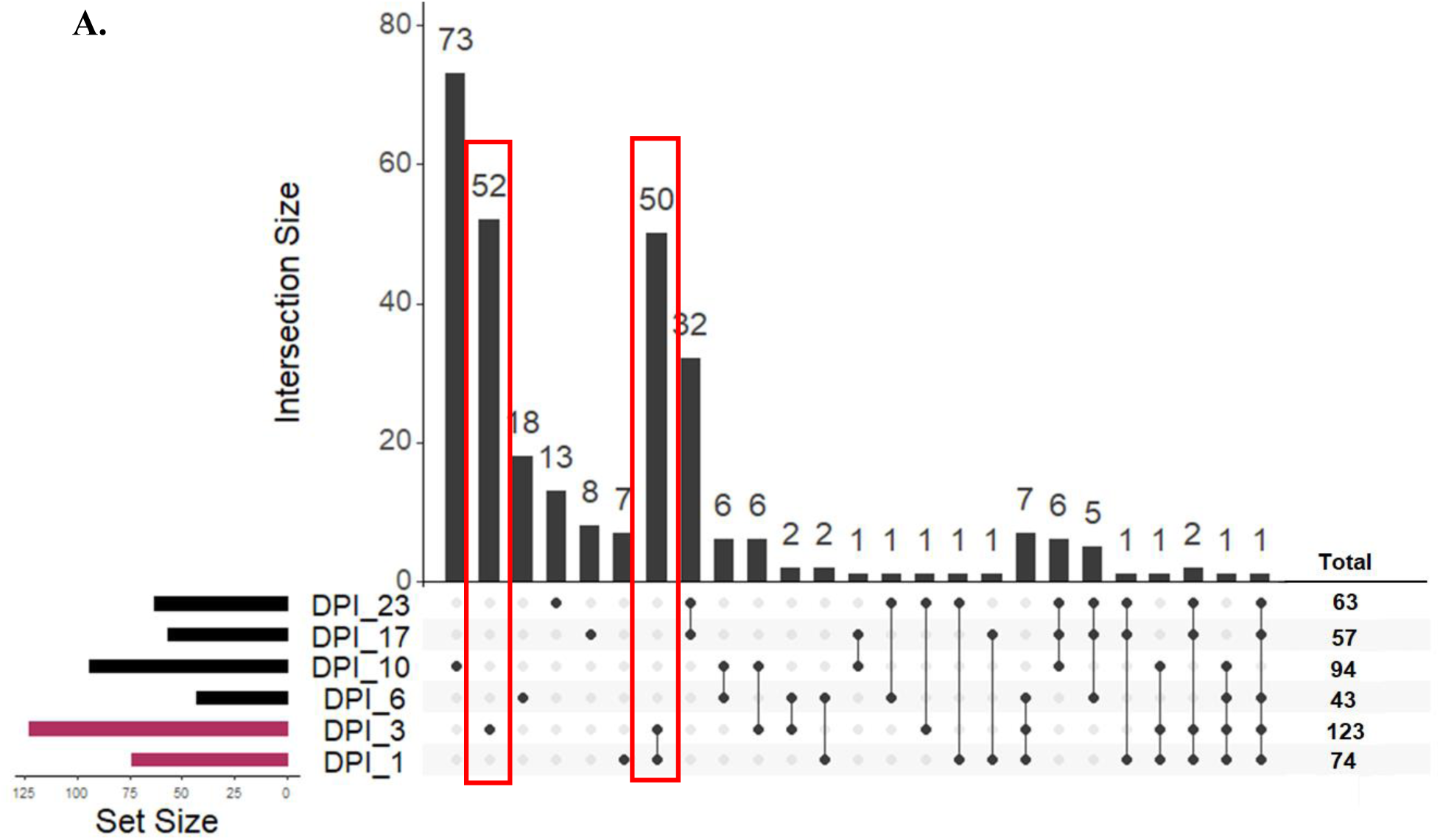
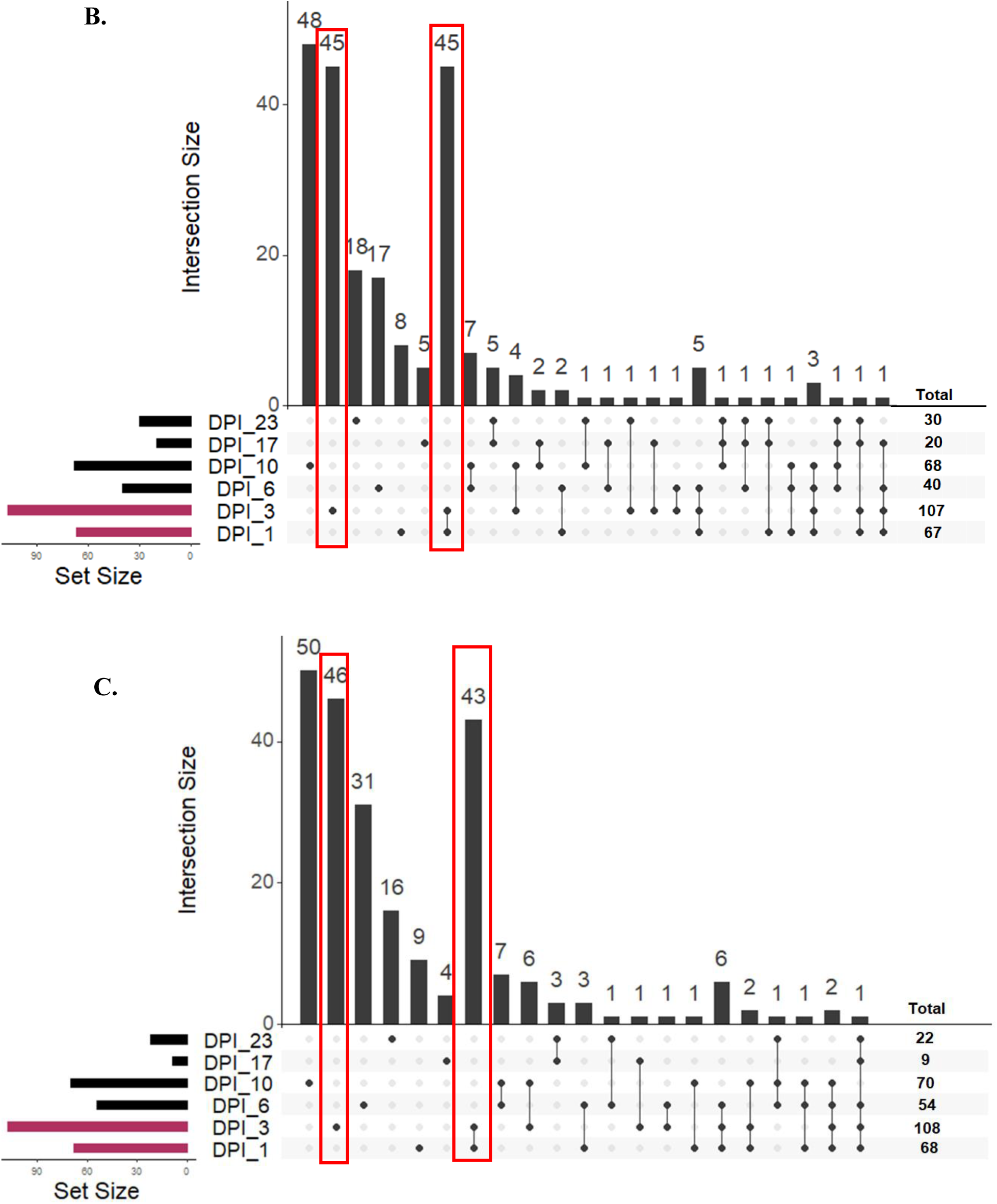
Numbers of unique and shared up-regulated differentially expressed genes (DEGs) of the wheat pathogen *Zymoseptoria tritici* among six time points during the: A) susceptible interaction on wheat cultivar Taichung 29; B) resistant interaction on the *Stb2-*carrying cultivar Veranopolis; and C) resistant interaction on the *Stb3*-carrying cultivar Israel 493compared to its non-host interaction on barley. The colored horizontal bars indicate the total number of up-regulated DEGs at the time points showing the highest number of uniquely up-regulated genes. The red rectangles highlight the numbers of genes that were exclusively differentially up-regulated at 1 DPI and at 1 and 3 DPI.

We also wanted to determine the unique and shared up-regulated DEGs at different sampling days during the resistant interaction on Israel 493 (*Stb3*) versus the non-host interaction on barley. In total, 46 up-regulated DEGs were exclusively observed at 3 DPI, followed by 43 up-regulated DEGs shared at 1 and 3 DPI (Fig. 7C). The peak of up-regulated genes at 3 DPI, during the early stage of infection, represents a higher number of up-regulated DEGs than those up-regulated at 10 DPI (Fig. 7C). This supports the idea of an early recognition of the host species from 1 to 3 DPI in the resistant interactions, and the subsequent activation of a large number of genes at 3 DPI. Presumably, the fungus recognizes the host species and activates a similar gene expression program during the first three days after inoculation on both susceptible and resistant hosts to initiate colonization of the plant leaves.

### Gene Ontology and enrichment analyses during the non-host interaction on barley

The significantly enriched GO categories among the down-regulated genes at 1 and 3 DPI in the comparisons of both susceptible and resistant interactions against the non-host interaction reveal some insights into the early response of the pathogen during the interaction on the non-host species. The enriched GO terms include those involved in the unfolded protein response (UPR) in fungi including unfolded protein binding (GO:0051082), and protein folding chaperone (GO:0044183) (Fig. 8). UPR is a stress-adaptive pathway that is activated under adverse environmental conditions and is involved in regulation of virulence traits in fungi (Guirao-Abad et al., 2022; Weichert et al., 2020).

**Fig. 8.**
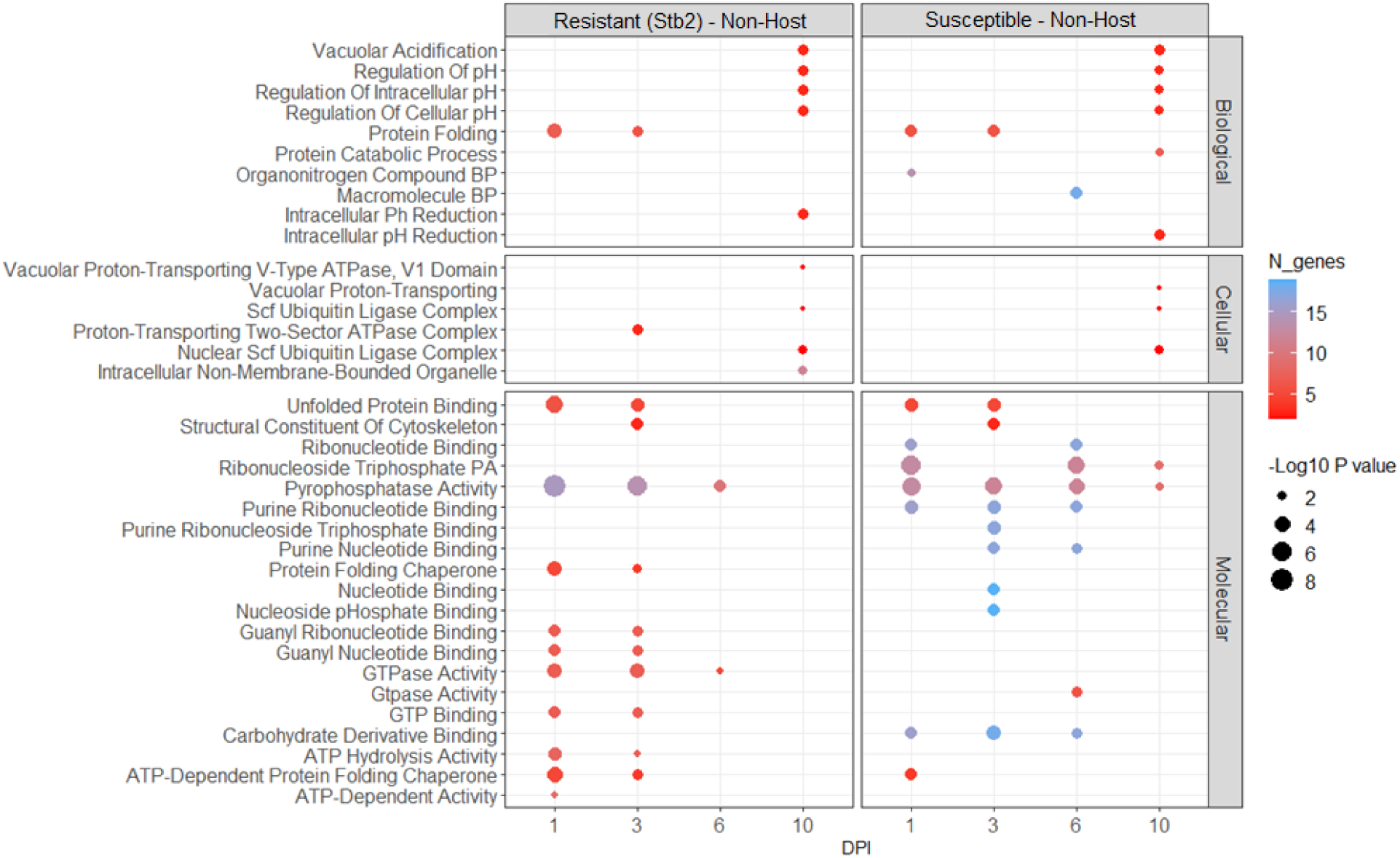
Gene Ontology terms enriched at 1, 3, 6 and 10 days post inoculation (DPI) with the wheat pathogen *Zymoseptoria tritici* in the non-host interaction on barley compared to the resistant interaction on the *Stb2-*containing cultivar Veranopolis (left panel) and on the susceptible wheat cultivar Taichung 29 (right panel). Only down-regulated genes (with the non-host interaction as the baseline) were included in the GO analysis. The number of down-regulated genes included in the GO analysis is 148 for the resistant interaction on the *Stb2*-containing cultivar Veranopolis compared to the non-host interaction at 1, 3, 6 and 10 DPI, and 145 genes for the susceptible interaction compared to the non-host interaction at 1, 3, 6 and 10 DPI. BP: Biosynthetic process.

Additionally, the down-regulated genes in *Z. tritici* at 10 DPI in the comparison between the susceptible interaction and the non-host interaction were significantly enriched in GO categories including vacuolar acidification (GO:0007035), intracellular pH reduction (GO:0051452), regulation of intracellular pH (GO:0051453), vacuolar proton transporting (GO:0000221), ubiquitin ligase complex (GO:0019005) and nuclear ubiquitin ligase complex (GO:0043224) (Fig. 8).

The KEGG pathway enrichment analysis showed that 5 pathways were significantly enriched among the up-regulated genes at 1 DPI during the susceptible interaction compared to the non-host interaction. These were glycolysis/gluconeogenesis, carbon metabolism, biosynthesis of secondary metabolites, biosynthesis of antibiotics and fructose and mannose metabolism. At 3 DPI three pathways were enriched, which include pyruvate metabolism, carbon metabolism and glycolysis and gluconeogenesis.

Three glycoside hydrolases (GHs) were significantly up-regulated at 1 DPI, and 6 CAZymes were up-regulated at 3 DPI, including 1 glycosyl transferase (GT), 4 GHs and 1 carbohydrate-binding molecule (CBM). At 6 and 10 DPI, 3 GHs, 5 GTs and 1 CBM were up-regulated (Supplementary Fig. S3). Also at 1 and 3 DPI, we were able to detect 7 and 9 SSPs that were up-regulated in the susceptible versus the non-host interaction.

### Expression of *Z. tritici* candidate effectors during early disease development

To gain further insight into *Z. tritici* pathogenicity, we used a set of established programs for effector identification (Hallgren et al., 2022; Sperschneider & Dodds, 2022; Teufel et al., 2022) using the predicted *Z. tritici* secretome, and selected thirty-one candidates with increased transcript abundance during the early stages of infection (1 and 3 DPI) in the susceptible interaction compared to the non-host interaction. The thirty-one selected effector candidates range in size from 68 to 554 amino acids (Table 4). Although the majority of the putative effectors do not contain subcellular-targeting peptide sequences, several encode predicted nuclear localization signals (NLS), chloroplast transit peptides (cTPs), or mitochondrial-targeting sequences (Table 4). One putative effector, Mycgr394290, includes multiple predicted subcellular localization signals including a cTP, mitochondrial targeting sequence, and a NLS (Table 4). Further examination of the candidate effectors revealed that two proteins, Mycgr3101991 and Mycgr3102792, contain predicted NLS, and one, Mycgr3106456, encoded a predicted cTP. Although most of the candidate effectors are annotated as uncharacterized, two are annotated as having peptidoglycan-binding LysM domains. Furthermore, two effector candidates encode Hce2-like domains, named after the homologs of *Cladosporium fulvum* Ecp2 (Stergiopoulos et al., 2012). Ecp2 is a secreted protein initially identified as a virulence factor in *C. fulvum* that belongs to a widely distributed superfamily of putative fungal effectors (Stergiopoulos et al., 2012). Another effector candidate is annotated as an allergen V5/Tpx-1-related protein, another as a heat shock protein, and one as a cerato-platanin protein. The putative function of members of these families as fungal effectors is described below.

**Table 4.** Characteristics of the 31 *Zymoseptoria tritici* candidate effectors that were selected for further analysis based on differential expression at the early or transition stages of the infection process.

| ID | Amino acids | Signal peptide | Protein size (kDa) | LOCALIZER prediction | Annotation |
| --- | --- | --- | --- | --- | --- |
| Mycgr394290 | 184 | 18-19 | 18.61 | Chloroplast (95-118)<br>Mitochondria (68-88)<br>NLS | Uncharacterized |
| Mycgr3111221 | 232 | 16-17 | 22.43 | —<sup>a</sup> | Peptidoglycan-binding LysM |
| Mycgr3106502 | 89 | 18-19 | 8.01 | — | Uncharacterized |
| Mycgr3104794 | 157 | 20-21 | 15.15 | — | Uncharacterized |
| Mycgr3109991 | 235 | 20-21 | 24.87 | NLS | Uncharacterized |
| Mycgr3105487 | 97 | 18-19 | 8.7 | — | Peptidoglycan-binding LysM |
| Mycgr3109710 | 200 | 26-27 | 18.8 | — | Allergen V5/Tpx-1 related |
| Mycgr3106125 | 70 | 23-24 | 4.95 | — | Uncharacterized |
| Mycgr3111223 | 554 | 18-19 | 58.81 | — | Heat shock protein DnaJ, N-terminal |
| Mycgr3110220 | 131 | 19-20 | 12.88 | — | Uncharacterized |
| Mycgr3107904 | 170 | 23-24 | 15.65 | — | Hce2 domain-containing protein |
| Mycgr346866 | 427 | 22-23 | 41.15 | — | Carbohydrate kinase, PfkB |
| Mycgr383064 | 75 | 18-19 | 6.15 | — | Uncharacterized |
| Mycgr3102792 | 114 | 19-20 | 10.63 | NLS | Uncharacterized |
| Mycgr379161 | 68 | 19-20 | 4.98 | — | Uncharacterized |
| Mycgr3106106 | 140 | 20-21 | 14.44 | — | Uncharacterized |
| Mycgr3111636 | 159 | 19-20 | 14.63 | — | Hce2 domain-containing protein |
| Mycgr3109137 | 154 | 18-19 | 15.81 | — | Uncharacterized |
| Mycgr3102996 | 163 | 23-24 | 15.12 | — | Uncharacterized |
| Mycgr3110887 | 283 | 19-20 | 30.59 | — | Uncharacterized |
| Mycgr3104009 | 483 | 25-26 | 51.39 | — | Beta-Ig-H3/fasciclin |
| Mycgr339947 | 149 | 17-18 | 13.95 | — | Cerato-platanin |
| Mycgr3105223 | 188 | 23-24 | 18.19 | — | Uncharacterized |
| Mycgr3105182 | 143 | 18-19 | 13.88 | — | Uncharacterized |
| Mycgr3104697 | 149 | 19-20 | 13.49 | — | Uncharacterized |
| Mycgr3101652 | 77 | 21-22 | 6.06 | — | Uncharacterized |
| Mycgr3106456 | 193 | 30-31 | 16.59 | Chloroplast (27-65) | GPI-anchored domain-containing protein |
| Mycgr3106329 | 135 | 19-20 | 13.06 | — | Uncharacterized |
| Mycgr382936 | 155 | 20-21 | 13.97 | — | Plastocyanin-like |
| Mycgr399161 | 164 | 15-16 | 16.03 | — | Uncharacterized |
| Mycgr3102849 | 285 | 20-21 | 27.95 | — | Uncharacterized |

<sup>a</sup> No specific prediction made.

Some of the effector candidates with annotations have been predicted to function as effectors or toxic compounds in other fungal plant pathogens, and some have been observed to be differentially expressed in previous transcriptome studies on *Z. tritici* (Mirzadi Gohari et al., 2015; Rudd et al., 2015; Yang et al., 2013). For example, five Hce2 domain-containing effectors have been identified in *Valsa mali*, the causal agent of Valsa canker in apple. One of these effectors induced cell death when transiently expressed in *N. benthamiana* (Zhang et al., 2019). In the genome of the wheat pathogen *Fusarium graminearum*, there are two genes that encode putative cerato-platanin proteins, which function as elicitors of defense responses on *Arabidopsis* leaves (Quarantin et al., 2019). A novel cerato-platanin-like effector protein from *Fusarium oxysporum* f. sp *cubense* triggers a hypersensitive response and systemic acquired resistance in tobacco (Li et al., 2019). Additionally, a *Sclerotinia sclerotiorum* cerato-platanin induces cell death in *N. benthamiana* and interacts with a PR-1 protein in the apoplast to facilitate infection (Yang et al., 2018). Finally, LysM domain-containing effectors from *Z. tritici* and *C. fulvum* have been previously characterized and shown to suppress chitin-induced plant immunity and to protect fungal hyphae from plant chitinase activity (Marshall et al., 2011; Sánchez-Vallet et al., 2013, 2020; Tian et al., 2021). Overall, the evidence from other fungal pathogens suggests that the candidate effectors identified in the present study could have a functional role at the early stages of infection.

The gene expression profiles of the thirty-one selected effector candidates were compared across all time points using the log-transformed normalized counts of each gene in the individual samples of the susceptible interaction. Our analyses reveal three clusters of candidate effector genes based on similarities in their gene expression patterns (Fig. 9). Each cluster contains candidate effector genes that are commonly regulated at specific time points. We found a cluster consisting of 10 candidate effectors that were commonly up-regulated at 1 and 3 DPI compared to the average relative expression of the 31 selected candidate effector genes. A second cluster, containing 10 commonly up-regulated genes, was identified at 10 and 17 DPI, and a third cluster of 11 genes was identified at 23 DPI (Fig. 9).

**Fig. 9.**
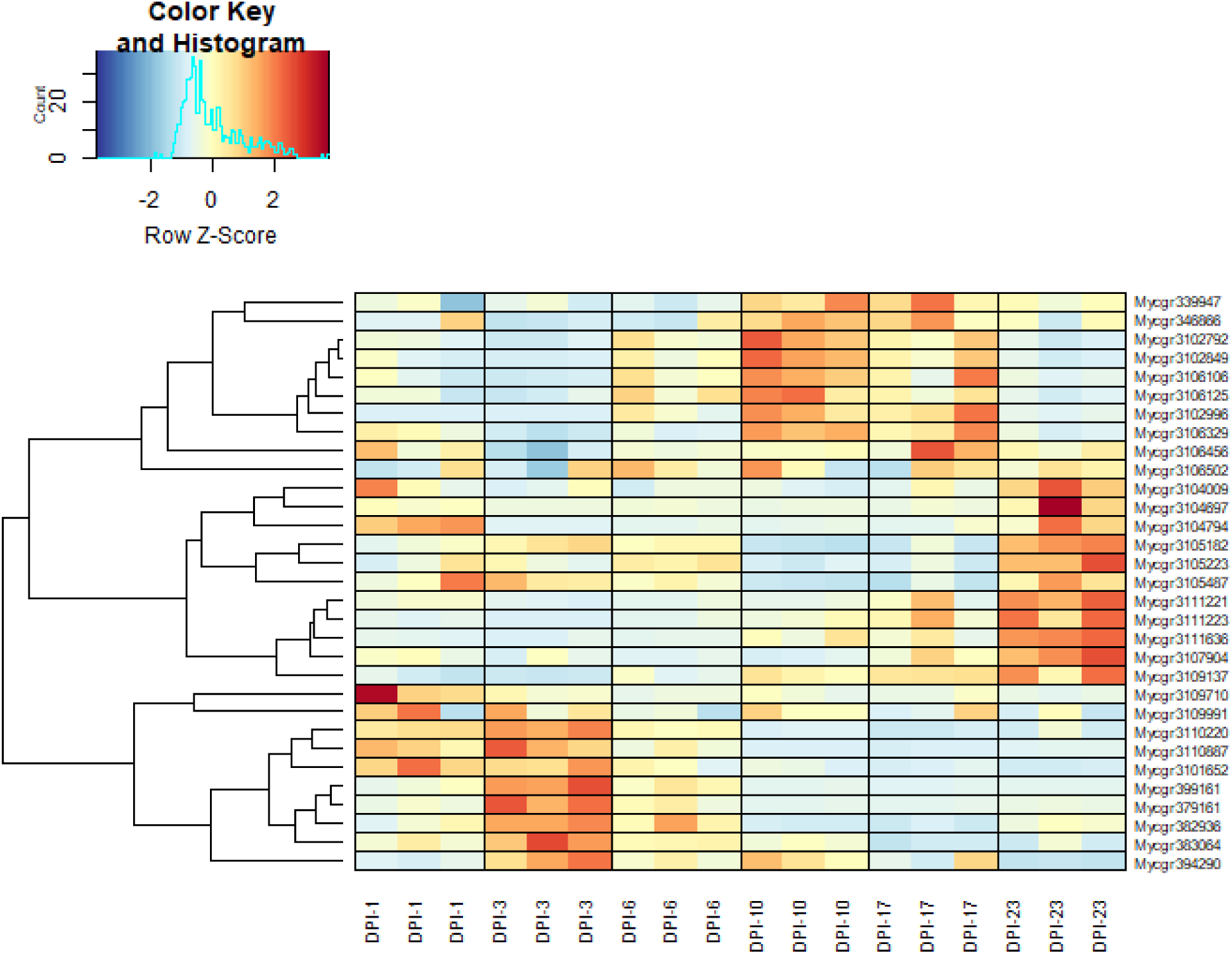
Relative expression of 31 selected *Zymoseptoria tritici* effector candidates over a 23-day time course in the susceptible wheat cultivar cultivar Taichung 29. The heat map shows the normalized expression value of each effector candidate in the individual samples for each time point for the susceptible interaction on cultivar Taichung 29 and not relative to another interaction. The dendrogram on the left shows the clustering of the candidate effectors based on their relative expression calculated by the Pearson correlation. All of these 31 effectors were expressed early when compared to the non-host interaction.

### Subcellular localization patterns of *Z. tritici* candidate effector-fluorescent protein fusions

Since several candidate effector proteins encode predicted subcellular-targeting sequences, we investigated the localization of six of these putative effectors to plant cellular compartments. These effectors were chosen based on their gene expression profiles (Fig. 9), with three (Mycgr3109710, Mycgr394290, Mycgr3106502) exhibiting higher expression at either 1 or 3 DPI, and the other three (Mycgr3107904, Mycgr3104794, Mycgr3111221) showing increased expression after 10 DPI. The localization of specific proteins can be determined by creating fluorescent protein (FP) fusion constructs (Berg & Beachy, 2008), where an FP is attached to the protein of interest, enabling visualization within intact tissues (Tanz et al., 2013). This approach has been applied to study viral and fungal proteins, allowing researchers to examine their interactions with plant organelles (Helm et al., 2022; Li et al., 2020; Petre et al., 2016). To test the localization of the six putative effectors, we fused super Yellow Fluorescent Protein (sYFP) to the C-terminus of candidate effector sequences, with and without their predicted signal peptides. We also examined the subcellular localization of the remaining 21 putative effectors using their mature protein sequences (lacking predicted signal peptides) and include the results in the supplementary data. These constructs were transiently expressed in *Nicotiana benthamiana* using *Agro-bacterium*-mediated transformation (agroinfiltration), and their localization patterns were examined using laser-scanning confocal microscopy (Helm et al., 2022; Jaiswal et al., 2023; Rogers et al., 2024). We used free sYFP as a negative control to establish a baseline for comparison, as it is a derivative of green fluorescent protein (GFP) and is used commonly to assess protein localization in plant cells (Berg & Beachy, 2008; Tanz et al., 2013). Due to their small size, FPs localize to both the cytoplasm and nucleus, providing a reference for identifying specific protein localization (Tanz et al., 2013).

In our analysis, if the fluorescence signal from an effector-sYFP fusion is observed in both the nucleus and cytosol, we consider the localization non-informative, as it matches the distribution of free sYFP. This suggests that the effector lacks a location-specific signal sequence to direct the fusion protein to a particular compartment. However, if the fluorescence signal is detected in a distinct organelle rather than both the nucleus and cytosol, so it is distinguishable from the localization of free sYFP, it indicates that the effector likely contains a targeting sequence that directs the sYFP to a specific subcellular compartment, confirming its potential localization.

Additionally, one challenge of using FPs in plants is the natural autofluorescence of cellular components such as cell walls and plastids, which can overlap with FP signals (Tanz et al., 2013). However, modern confocal microscopes can largely account for this background auto-fluorescence (Tanz et al., 2013). To further rule out autofluorescence, we examined *N. benthamiana* leaves infiltrated with *Agrobacterium* expressing a protein construct tagged with hemagglutinin (HA) instead of sYFP (Supplementary Fig. S5) and also imaged an uninfiltrated leaf at the sYFP excitation wavelength under the confocal microscope (Supplementary Fig. S6). In both cases we did not observe autofluorescence.

Live-cell imaging of *N. benthamiana* epidermal cells revealed that fluorescence signals from Mycgr3111221, Mycgr3106502, and Mycgr3104794, when expressed without their signal peptides, were indistinguishable from those of the free sYFP controls and were thus considered to have a non-informative localization (Fig. 10A). The fluorescence signal from Mycgr3107904:sYFP without its signal peptide was predominantly observed in the cytosol (Fig. 10A), suggesting that Mycgr3107904 may be cytosolic. However, it also could be associated with the plasma membrane or the apoplastic space, and further analyses will be required to confirm its precise localization. Intriguingly, Mycgr3109710:sYFP and Mycgr394290:sYFP, when expressed without their signal peptides, labeled what appear to be cytosolic bodies or punctuate structures, and these punctuate structures appear to be mobile for Mycgr3109710:sYFP (Fig. 12, Supplementary video 1).

**Fig. 10.**
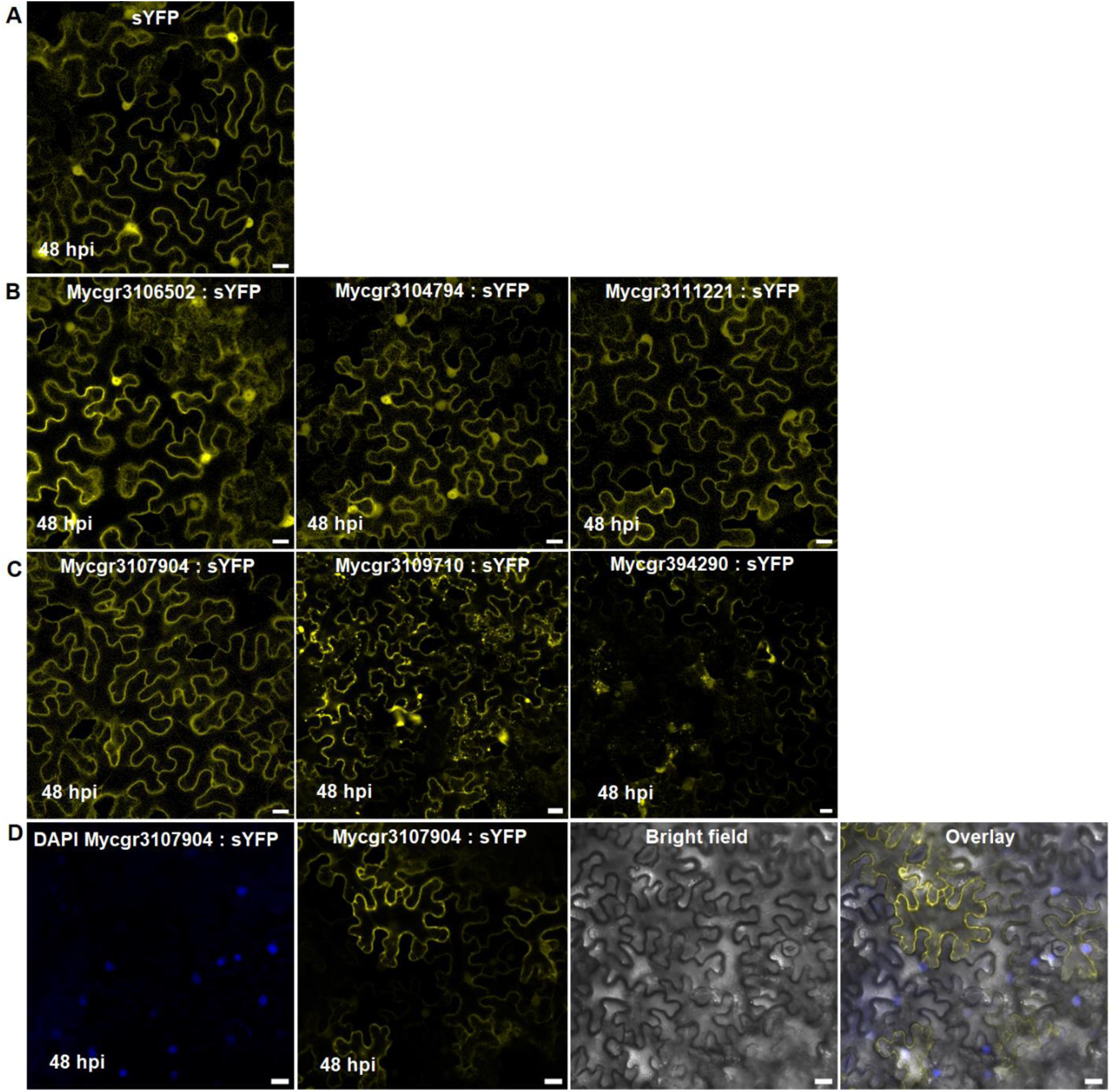
Subcellular localization of six *Zymoseptoria tritici* effectors in *Nicotiana benthamiana* leaf cells expressed without their putative signal peptides. A) Free sYFP localizes to the nucleus and cytosol of the cell, and was included as a negative control to establish a baseline for comparison; B) Z*ymoseptoria tritici* candidate effectors Mycgr3106502, Mycgr3104794 and Mycgr3111221 display a non-informative localization that is indistinguishable from the localization of the negative control free sYFP; C) Mycgr3107904 localizes only to the cytosol and no localization signal is detected in the nucleus; Mycgr3109710- and Mycgr394290-fluorescent proteins accumulate in what appears to be mobile (Supplementary video) cytosolic elements in *N. benthamiana* cells. D) DAPI nucleus staining is shown as a nuclear marker for comparison using effector Mycgr3107904 that localizes predominantly to the cytosol. sYFP-tagged *Z. tritici* candidate effectors were transiently expressed without their putative signal peptides in *N. benthamiana* leaves using agroinfiltration. Confocal micrographs of leaf epidermal cells were captured 48 hours following agroinfiltration. The scale bars shown at the bottom right of each image represent 20 µm. All confocal micrographs are of single optical sections.

**Fig. 11.**
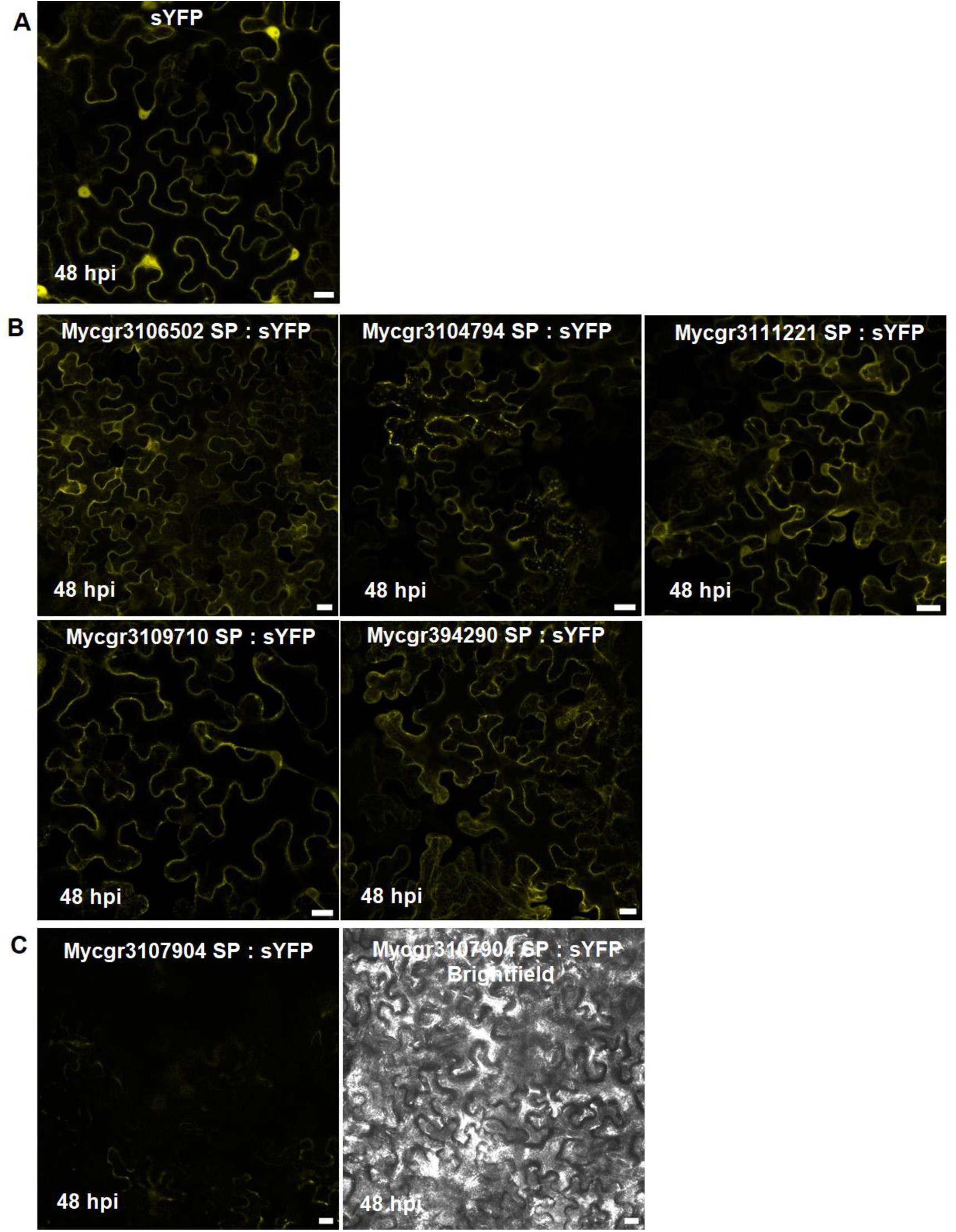
Subcellular localization of six *Zymoseptoria tritici* effectors in *Nicotiana benthamiana* leaf cells expressed with their putative signal peptides. A) Free sYFP localizes to the nucleus and cytosol of the cell, and was included as a negative control to establish a baseline for comparison; B) Z*ymoseptoria tritici* candidate effectors were expressed with their putative signal peptides. Mycgr3106502, Mycgr3104794, Mycgr3109710, and Mycgr394290 localize predominantly to the cytosol and no localization signal is detected in the nucleus. Mycgr3111221 localizes to both the cytosol and nucleus; C) No fluorescence signal was detected for Mycgr3107904. The bright-field section for this effector is included to show the absence of the fluorescence signal. sYFP-tagged *Z. tritici* candidate effectors were transiently expressed with their putative signal peptides in *N. benthamiana* leaves using agroinfiltration. Confocal micrographs of leaf epidermal cells were captured 48 hours following agroinfiltration. The scale bars shown at the bottom right of each image represent 20 µm. All confocal micrographs are of single optical sections.

When expressed with their native signal peptides, Mycgr3106502, Mycgr3104794, Mycgr3109710, and Mycgr394290 predominantly localized to the cytosol, with no detectable fluorescence in the nucleus (Fig. 10B). This distinct pattern, compared to the nucleus-cytosol localization of the free sYFP control, suggests that these effectors may be cytosolic. However, further observations including co-localization markers and plasmolysis are required to exclude the potential localization of these effectors to the plasma membrane or the apoplastic space. In contrast, Mycgr3111221, when expressed with its signal peptide, labeled both the cytosol and nucleus, resulting in a non-informative localization pattern indistinguishable from that of free sYFP. Mycgr3107904 did not show a fluorescent signal at 48 hours post infiltration, likely indicating protein degradation or rapid delivery into the apoplast (Fig. 10B).

Immunoblot analyses were used to confirm the expression and integrity of the *Z. tritici* effector–fluorescent protein fusions in *N. benthamiana*. This assay revealed that, of the twenty-seven candidate effectors screened, twenty-six accumulated at detectable levels when transiently expressed without their predicted signal peptides (Supplementary Fig. S8). When expressed with their signal peptides, five of six predicted effectors showed detectable accumulation (Supplementary Fig. S9). While Mycgr3104794 accumulated at a low level, Mycgr3107904 did not produce an observable band in the immunoblot assay (Supplementary Fig. S9). Notably, for several effectors expressed with or without their signal peptides, we observed multiple bands in the immunoblots, often with one band corresponding to the expected size and additional bands of lower molecular weight (Supplementary Figs. S8 and S9). This pattern suggests possible post-translational cleavage, partial degradation, or the presence of processing intermediates, which may affect the stability or integrity of the effector fusions. In some cases, the lower-molecular-weight bands were approximately the size of free sYFP, raising the possibility that the fluorescence observed in both the cytosol and nucleus for some effectors may be due to free sYFP rather than the intact fusion protein.

## Discussion

Comparative RNAseq analysis clearly shows that different suites of genes are expressed by *Z. tritici* during its biotrophic and necrotrophic infection stages. The comparison of the infection time course in the susceptible host and the two resistant hosts, allows us to identify genes that are necessary for or participate in the transition from the biotrophic to the necrotrophic growth stages. The sudden increase in the number of differentially expressed genes at 10 days post inoculation coincides with the onset of visible lesions on the leaves (Supplementary Fig. S1). This aligns with expectations, as previous studies have demonstrated that *Z. tritici* undergoes significant transcriptomic reprogramming during the transition to its necrotrophic phase (Kellner et al., 2014; Palma-Guerrero et al., 2017; Rudd et al., 2015; Yang et al., 2013) prior to the exponential growth in fungal biomass that occurs in susceptible cultivars beginning 18-20 DPI (Adhikari et al., 2025). Our results provide the first confirmation that the main difference in the pathogen gene expression in a susceptible compared to a resistant interaction occurs at the transition stage rather than during the early infection stage (1-6 DPI), consistent with the low levels of fungal biomass reported in both resistant and susceptible interactions before 10 DPI (Adhikari et al., 2025). These results also reveal that *Z. tritici* deploys a similar gene activation pattern and comparable molecular mechanisms during the symptomless early stage of the disease in a susceptible interaction and in resistant host interactions.

In previous comparisons of host and non-host interactions, investigators found that when other grass species, such as *Brachypodium distachyon* or *Agropyron repens* are inoculated with *Z. tritici*, the pathogen penetrates the leaf surface through stomata, but the infection is halted in the substomatal cavity (Kellner et al., 2014). This suggests an early recognition and interaction between the host and pathogen, during which *Z. tritici* can escape detection or manipulate host defenses in susceptible wheat lines to penetrate the leaves, colonize the plant, and initiate the symptomless phase (Kellner et al., 2014).

Comparison of gene expression in susceptible and resistant wheat cultivars to gene expression in a non-host allows us to examine genes involved in initiating/supporting host colonization and avoiding pathogen recognition. We suggest that some of these genes may play a role in early epiphytic growth, stomatal detection and penetration by *Z. tritici* during its biotrophic phase. Previous research shows that while *Z. tritici* can colonize epiphytically in susceptible, resistant, and some non-host species, its penetration attempt efficiency is significantly lower in barley accessions (about 10% less) compared to near-isogenic wheat lines (NILs) containing *Stb* resistance genes (Battache et al., 2024). This difference may help explain the variation in early-stage transcriptomic responses in host and non-host interactions. It is possible that *Z. tritici* possesses a more efficient mechanism for stomatal recognition and penetration of its host species, which could influence gene expression, survival, and pathogenicity.

Battache et al. (2024) found no difference in penetration attempt efficiency and a similar percentage of reached stomata, in susceptible and resistant interactions in NILs. Additionally, *Z. tritici* can undergo both asexual and sexual reproduction during its phase of epiphytic growth, even on resistant wheat cultivars (Fones et al., 2023). This is supported by the similar expression profiles that we observe in susceptible and resistant interactions from 1 to 6 DPI. This is particularly interesting, as previous research also has shown that *Z. tritici* can grow epiphytically for up to 10 days on both wheat and non-host species like barley (Battache et al., 2024; Fones et al., 2017). However, the efficiency of preventing stomatal penetration by *Z. tritici* varies according to the resistance types, with *Stb6*-mediated and barley non-host resistance being the most effective among those tested (Battache et al., 2024). Understanding the expression profile of *Z. tritici* associated with its extended epiphytic growth is an important area for future research which could provide valuable insights into the symptomless infection phase and help inform early-stage disease management strategies.

Because we noticed similar expression profiles in *Z. tritici* during both susceptible and resistant interactions at early time points, and given the resistance response in cultivars carrying *Stb6*, we became interested in the expression dynamics of *AvrStb6*, the apoplastic effector of *Z. tritici* that is recognized by Stb6. To explore this, we re-analyzed our RNAseq data (see Supplementary Excel material). Given that *AvrStb6* was not predicted in the JGI annotation (Goodwin et al., 2011) originally used for our analysis, we mapped the reads against a recently released *Z. tritici* genome annotation that correctly includes this gene (Lapalu et al., 2025). At 1 DPI, *AvrStb6* showed high expression in all host interactions (susceptible and resistant) compared to the non-host, where no transcripts were detected. Notably, expression was strongest in the resistant interaction with Israel493, while levels in the susceptible interaction and in Veranopolis were lower. At 3 DPI, *AvrStb6* expression decreased in all host interactions and remained absent in the non-host. By 10 DPI, expression was strongly upregulated in the susceptible interaction, coinciding with the appearance of the first visible disease symptoms. In contrast, both resistant interactions showed only low transcript counts at this stage, which likely reflects reduced fungal biomass in resistant plants. Across these time points, *AvrStb6* was never detected in the non-host interaction. These results suggest that *AvrStb6* is transiently expressed early during infection and peaks later in the susceptible interaction, supporting a role in virulence, while its recognition by *Stb6* explains its avirulence function in resistant cultivars.

Our results confirm that the transcriptomic response in *Z. tritici* is significantly different during the early infection stage in both susceptible and resistant interactions on the host species compared to the non-host interaction with barley. *Z. tritici* activates a different gene expression program, implying that it uses different molecular mechanisms at 1 and 3 DPI when interacting with susceptible or resistant wheat cultivars, compared to its program when interacting with a non-host species. This suggests that non-host species may recognize and neutralize *Z. tritici* at a very early stage, preventing the implementation in the pathogen of its normal biotrophic growth program. Future research on the mechanisms of host recognition through the analysis of non-host interactions should incorporate transcriptomic data with increased sequencing depth to capture low-abundance transcripts, especially during the early stages of disease and in non-host interactions where there is limited fungal growth. This deeper analysis should be complemented by additional experiments that accurately monitor fungal growth, development and plant colonization during non-host interactions. Such an integrated approach could provide insights into the mechanisms of host recognition by the pathogen and inform strategies for developing wheat cultivars with broad-spectrum resistance.

The highest number of DEGs in *Z. tritici* when comparing the susceptible interaction on the cultivar Taichung 29 with both resistant interactions on Veranopolis and Israel 493 is seen at 10 DPI. Interestingly, based on a previous principal component analysis (PCA), Yang et al. (2013) identified 13 DPI as the point with the greatest difference in transcription in a susceptible interaction with the wheat cultivar Sevin. However, inspection of their Figure 3 shows that the 10 DPI and 13 DPI samples are very similar and that the largest change is actually between 4 DPI and 10 DPI. This supports our finding that the most important transcriptome reprogramming in the pathogen, in a susceptible compared to a resistant interaction, occurs during the biotrophic/necrotrophic transition stage (10 DPI) rather than during the necrotrophic phase (13 DPI), revealing molecular mechanisms in *Z. tritici* that are activated at 10 DPI and determine the outcome of the infection in each type of interaction. Rudd et al. (2015) found that, in a susceptible interaction compared to growth in Czapek-Dox broth culture medium, the highest number of up-regulated genes in *Z. tritici* was observed during the transition to the necrotrophic stage (9 DPI), while in our study, it occurred at 10 DPI. Rudd et al. (2015) identified 1,389 DEGs at 9 DPI, while we found 601 and 587 total DEGs in the susceptible interaction compared to the resistant *Stb2*-gene and *Stb3*-gene interactions, respectively. Again, these findings indicate that significant transcriptional reprogramming in *Z. tritici* occurs during the transition stage, around 9-13 dpi, in a susceptible interaction compared to fungal growth in culture media or to infection of a resistant host.

Yang et al. (2013) demonstrated distinct transcriptional responses during different phases of the infection. Their PCA showed separation between the 4-day, 10-day, and 13-day infected samples, representing the asymptomatic phase, the intermediate transition stage, and the necrotrophic phase, respectively. In our analyses, PCA identified four clusters for the replicates of the susceptible interaction (C1: 1 and 3 DPI, indicated by the red circles and diamonds on Fig. 1; C2: 6 DPI, denoted by the red triangles on Fig. 1; C3: 10 DPI, red squares; C4: 17 and 23 DPI, red circles and diamonds on Fig. 2). C1 contains all the replicates from both susceptible and resistant host interactions, revealing a similar gene activation in the fungus at 1 and 3 DPI. From 6 DPI, we observe the beginning of a change in transcriptional response in the susceptible and resistant interactions according to PC2 of the PCA (Fig. 1). This time point marks the separation of replicates from the susceptible interaction into their own cluster. In C3 and C4 gene expression patterns in the susceptible interactions are clearly separated from all replicates of the resistant interactions (Figs. 1 and 2). This finding strongly suggests the existence of defining molecular processes in the pathogen that govern or result from the successful switch from the biotrophic phase to the necrotrophic stage at 10 DPI, ultimately determining whether the interaction is susceptible or resistant. In our analyses, 1,504 genes at 1, 3, 6, and 10 DPI and 222 genes at 17 and 23 DPI were detectable and passed the normalization and filtering criteria for the comparison of the susceptible and resistant interactions. It is worth noting that DEGs identified at 10 DPI account for approximately 40% of the normalized read counts, highlighting that expression of a large fraction of all genes is reprogrammed during the transition to the necrotrophic stage in the susceptible interaction. These genes are likely involved in molecular pathways that determine the ability of the pathogen to overcome the plant defense response and induce symptoms at around 10 DPI in a susceptible interaction. Adhikari et al. (2025) showed that fungal biomass in two susceptible interactions begins to increase exponentially after 18 DPI. The transcriptional changes indicated by the clear separation of C4 in Fig. 2 reflects and supports this exponential growth of fungal biomass. Interestingly, as demonstrated by previous research, changes in gene expression of the pathogen are mirrored by those in the host at early and late points in the infection process (Adhikari et al., 2007). The host responds during early detection of the pathogen and then again when the pathogen transitions from biotrophic to necrotrophic growth (Adhikari et al., 2007).

KEGG analysis identified biosynthesis of antibiotics and biosynthesis of secondary metabolites as significantly enriched pathways in the susceptible interaction compared with both resistant interactions at 10 DPI. Similarly, in a study conducted on *Bipolaris maydis* on resistant and susceptible non-CMS (cytoplasmic male sterile) maize genotypes, these same biosynthetic pathways were exclusively enriched in susceptible interactions (Meshram et al., 2022). The authors concluded that the successful pathogenicity of *B. maydis* depends on pathogen genes such as toxin-related effector molecules, cell wall degradation of the host, detoxification, and host defense evasion (Meshram et al., 2022). These findings agree with our observation that *Z. tritici* activates an arsenal of biosynthetic pathways, CAZymes including CWDEs, and SSPs during the transition to the necrotrophic stage in the compatible interaction. However, among the down-regulated genes, although we could not find significantly enriched pathways due to the low number of genes that were annotated, we identified down-regulated CAZymes and SSPs at 10 DPI, such as the alcohol dehydrogenase CAZyme family (AA3_2). This group includes enzymes with important roles in the fungal life cycle and colonization of the host (Gutiérrez-Corona et al., 2023). In *B. cinerea,* an alcohol dehydrogenase regulates fungal development, environmental adaptation and pathogenicity as observed in the morphology and virulence of knock-out mutants (DafaAlla et al., 2022). The benzoquinone reductase family (AA6) also was down-regulated, which contains intracellular enzymes involved in the biodegradation of aromatic compounds and in the protection of fungal cells from reactive quinone compounds (Levasseur et al., 2013).

In the comparison of the susceptible and the resistant interactions to the non-host interaction with barley, the main difference in the *Z. tritici* transcriptional responses occurs at 1 and 3 DPI in all the comparisons evaluated. We identified 285, 261 and 269 DEGs at 1 and 3 DPI in the comparisons of the susceptible, the resistant *Stb2*-gene, and the resistant *Stb3*-gene interactions with the non-host interaction, respectively. At 3 DPI the numbers of DEGs were always higher than those at 10 DPI for the three comparisons, which reveals that the main difference in the pattern of gene expression in the fungus during a host interaction compared to a non-host interaction occurs at the early stage of infection rather than during the transition stage. We detected 352 genes with measurable counts at 10 DPI, indicating that the pathogen is able to grow and colonize to a certain extent in the non-host species. In a previous report (Kellner et al., 2014), the patterns of transcription in *Z. tritici* during infection in the susceptible wheat cultivar Obelisk and in the non-host grass species *B. distachyon* were evaluated after 4 DPI. They found that genes encoding proteins with putative secretion signals were enriched among those up-regulated during infection of wheat and genes with plant-specific expression patterns. In our analysis, we observe a higher proportion of small, secreted proteins (SSPs) in the up-regulated sets of genes at 1 and 3 DPI compared to the down-regulated sets.

Down-regulated genes at 1 and 3 DPI in the host interactions, compared to the non-host interaction, were enriched in GO terms linked to the unfolded protein response (UPR). UPR is a stress-adaptive pathway that modulates the accumulation of misfolded proteins in the endoplasmic reticulum (ER), typically activated when the pathogen faces an increased demand for protein secretion (Weichert et al., 2020). It also helps pathogenic fungi survive under adverse environmental conditions, such as the stress of infection (Guirao-Abad et al., 2022). Notably, plant-pathogenic fungi can exploit UPR to regulate virulence traits like thermotolerance, nutrient acquisition, effector secretion, and resistance to antimicrobial peptides (Weichert et al., 2020). In *Z. tritici*, the need to deploy more secreted proteins for effective epiphytic colonization, combined with a potential early resistance response from the non-host interaction, may create stress that triggers UPR activation to meet the demands for survival in such a challenging environment. Interestingly, at 10 DPI, genes down-regulated in the host interaction compared to the non-host were enriched for GO terms related to intracellular pH regulation and vacuolar acidification. Although intracellular pH in fungi is typically stable, it can shift rapidly in response to environmental stress, such as changes in extracellular pH or nutrient availability (Fernandes et al., 2017). In *Aspergillus niger*, extracellular acidification induces a transient drop in cytosolic pH. Similar responses in *Fusarium oxysporum* and *Saccharomyces cerevisiae* lead to MAPK reprogramming, suggesting a conserved role for pH-dependent signaling in morphogenesis and invasion (Fernandes et al., 2017; 2023). Acid stress has also been shown to promote fungal survival by triggering growth arrest (Lucena et al., 2020), while heat shock and cell wall stress cause transient intracellular acidification that enhances stress tolerance (Triandafillou et al., 2020; Garcia et al., 2017). The upregulation of genes involved in vacuolar acidification and pH regulation during the non-host interaction suggests that *Z. tritici* may be activating adaptive mechanisms to counteract stress, maintain pH homeostasis, and enable survival under unfavorable conditions.

Among the 31 selected effector candidates, an interesting finding is that effector candidate Mycgr3109710 localized to mobile cytosolic bodies near the cell periphery when expressed without its signal peptide in *N. benthamiana*, whereas with its signal peptide, it was restricted to the cytosol. Immunoblot analysis showed a clear band corresponding to the expected size of the fusion protein in both cases, along with a lower band corresponding to free sYFP, suggesting some degree of degradation or cleavage (Supplementary Fig. S8). However, the observed fluorescence signal did not follow the typical diffuse nucleus-cytosol pattern associated with free sYFP, indicating that the effector is likely localizing to specific compartments. This effector is annotated as part of the Allergen V5/Tpx-1-related group, containing CAP/SCP domains characteristic of the PR-1-like superfamily (Wangorsch et al., 2022). BLAST analysis confirmed homology with PR-1-like proteins from *Zymoseptoria brevis* and *Aureobasidium* sp. While PR-1-like proteins are well studied in plants and animals, few studies exist in filamentous ascomycetes. Notably, in *Fusarium graminearum*, deletion of a PR-1-like gene reduced virulence (Lu & Edwards, 2018), and in *Moniliophthora* species, these genes are upregulated during biotrophic stages (Teixeira et al., 2012). Mycgr3109710 also shows early infection expression (1 and 3 DPI), supporting its potential role in host–pathogen interactions. We hypothesize that Mycgr3109710 targets the plant vesicle-trafficking system, given its localization to mobile punctate structures resembling the multivesicular body localization reported for PR-1 variants from *Arabidopsis thaliana* when expressed in *N. benthamiana* (Pečenková et al., 2022). To further confirm the localization of Mycgr3109710 to mobile cytosolic bodies in *N. benthamiana*, we recorded a time-lapse series capturing the localization of the effector. This analysis revealed the dynamic motility of the observed cytosolic elements (Supplementary video 1).

Similarly, Mycgr394290, when expressed without its signal peptide, localizes to the nucleus and labels cytosolic punctate structures with less mobility than those labeled by Mycgr3109710. When expressed with its signal peptide, Mycgr394290 localizes predominantly to the cytosol. We suggest that Mycgr394290, without its signal peptide, may target organelles involved in sorting and trafficking of molecules. An example of a fungal effector with similar localization is Foa3 from *Fusarium oxysporum*, which functions within the plant cell to suppress host defenses. Foa3 showed a weak signal in the cytosol but was strongly detected in punctate structures when a secretory signal peptide was included (Tintor et al., 2022), opposite to what we observed for Mycgr394290. However, their study demonstrates that Foa3 suppresses pattern-triggered defense responses to the same extent regardless of whether the protein carries the signal peptide (Tintor et al., 2022). Regarding the identity of the punctuate structures, Tintor et al. (2022) proposed that the labeled structures may represent vesicles containing proteins targeted for secretion. This is intriguing because Mycgr394290 encodes predicted mitochondrial, chloroplast, and nuclear localization signals according to LOCALIZER (Table 4). Although the exact identity of the punctate subcellular structures labeled by Mycgr394290:sYFP is uncertain, it is plausible that this protein acts to suppress the defense mechanisms of the plant in the same way as Foa3, which is known to suppress the plant immune response.

Another candidate effector, Mycgr3107904, was mainly localized to the cytosol only when expressed without its signal peptide. Interestingly, no fluorescent signal was detected when the effector was expressed with its signal peptide, likely suggesting degradation or rapid translocation into the apoplastic space. This effector is annotated as a Hce2 domain-containing protein. Hce2 (named from **h**omologs of ***C****. fulvum* **E**cp2 effector) is a domain that causes plant cell death, and is widely conserved in the fungal kingdom (Laugé et al., 1997; Zhang et al., 2019). There are three predicted Hce2-containing effectors in the genome of *Z. tritici*: MgEcp2 (Mycgr3104404); MgEcp2-2 (Mycgr3107904); and MgEcp2-3 (Mycgr3111636) (Stergiopoulos et al., 2010, 2012). In addition, a fourth homolog of Ecp2 was found to be up-regulated in *Z. tritici* at 8 DPI and then down-regulated at 12 DPI. This effector candidate induces necrosis in wheat in a cultivar-dependent manner (Ben M’barek et al., 2015). In another study (Mirzadi Gohari et al., 2015), Mycgr3104404, Mycgr3107904 and Mycgr3111636 were up-regulated during the transition to necrotrophy, and in the necrotrophic stage at 8 and at 12 DPI. In our study, Mycgr3111636 was up-regulated at 1 DPI and at 10 DPI but we were unable to transform this effector candidate into *Agrobacterium*, suggesting a possible toxic effect on *Agrobacterium* cells. We transformed Mycgr3107904, which was up-regulated at 10 DPI, and we found that the fluorescence signal was mainly localized to the cytosol. This is very interesting as the Ecp effector of *C. fulvum* is one of the few proteins that can induce necrosis in tomato and tobacco plants irrespective of the signal peptide, which indicates that its host target is intracellular (Ben M’barek et al., 2015). We did not observe fluorescence in a specific intracellular compartment, but the protein is localized within the cytosol, which suggests a plausible intracellular target protein in the plant.

In our analysis, Mycgr3111221, which is a Lysin (LysM) domain-containing effector, was up-regulated at the early stage of infection in the susceptible compared to the non-host interaction. There are three LysM effector genes identified and characterized in the *Z. tritici* genome (Marshall et al., 2011; Sánchez-Vallet et al., 2020; Tian et al., 2021). Two of them, Mycgr3111221 and Mycgr3105487, were previously shown to be up-regulated during the asymptomatic phase (Marshall et al., 2011; Tian et al., 2021). Interestingly, Mycgr3111221 suppresses chitin-induced plant immunity and protects fungal hyphae against plant chitinase activity (Marshall et al., 2011). Mycgr3111221 is a homolog of Ecp6 effectors in *C. fulvum,* which are secreted directly into the apoplastic space where they perform their chitin-binding function (Sánchez-Vallet et al., 2020). However, in our analysis, Mycgr3111221 shows a non-informative localization in both the nucleus and cytosol, which is indistinguishable from the localization of the free sYFP control. This occurs irrespective of the presence or absence of the putative signal peptide.

Our findings also underscore the limitations of using sYFP fusions to assess effector subcellular localization. While our localization assays provide some insight into the potential intracellular compartments targeted by *Z. tritici* candidate effectors, several caveats must be considered. Fusion of sYFP to the C-terminus may interfere with proper protein folding, stability, trafficking, or function, potentially resulting in mislocalization or loss of activity (Tian et al., 2004; Snapp, 2005). In addition, fluorescent proteins like sYFP contain protease-sensitive regions that may be cleaved during expression in plant cells, releasing the free fluorophore and producing misleading localization signals unrelated to the effector itself (Chen et al., 2013).

These limitations are supported by our immunoblot results, which reveal degradation products for several constructs. This is particularly evident for the candidate effectors expressed with their native signal peptides (Supplementary Fig. S9). The inclusion of the fungal signal peptide likely directs the protein into the plant secretory pathway, targeting it for secretion into the apoplast or extracellular space (Petre & Kamoun, 2014). Within this pathway, the fusion protein may undergo post-translational processing, including signal peptide removal or additional proteolytic cleavage at internal processing sites, as well as misfolding or degradation, especially in the endoplasmic reticulum or apoplast. Such proteolytic processing can result in activation, inactivation, and completely altered protein function (Rogers and Overall, 2013). These cleavage events could generate truncated protein fragments that are still detected by immunoblot but would not necessarily retain the fluorescent tag, potentially leading to misleading or incomplete localization patterns.

One type of cleavage site that could influence post processing is KEX2. Bioinformatics identification of KEX2 cleavage sites in secreted proteins from *Z. tritici* could provide insights into the proteolytic processing that some candidate effectors undergo within the secretory pathway, but so far this has not been performed. The KEX2 cleavage site is highly conserved among fungi and its characteristic motifs are recognized by Kex2 proteases involved in protein processing (Ding et al., 2016; Le Marquer et al., 2019). Studies on KEX2-processed repeat proteins (KEPs) in fungal secretomes revealed that plant pathogens displayed a larger number of KEPs than animal pathogens or saprotrophs (Le Marquer et al., 2019). They also found that, among fungal species with the largest numbers of KEPs, twelve of them interacted with plants either as symbionts, endophytes or pathogens, suggesting that KEPs may play a role in the diversity of mechanisms deployed by fungi to interact with their plant hosts (Le Marquer et al., 2019). Whether KEPs are important for *Z. tritici* is not known but their identification and analysis could be a subject for future research.

Additionally, because candidate effectors are secreted, they may not be present in the total intracellular protein extract, resulting in weak or undetectable signal in standard immunoblot analyses. Therefore, observed localization patterns should be interpreted with caution and ideally validated through complementary approaches, such as apoplastic fluid extraction, plasmolysis, confocal imaging with co-localization markers, or ER retention constructs. In this context, the immunoblot analyses presented in Supplementary Figs. 8-9 provide evidence of post-translational cleavage and degradation, and should be considered a primary line of support for the limitations of the localization experiments.

Future studies also should assess the biological relevance of these effectors by evaluating their potential roles in suppressing plant immunity through functional assays, including ROS burst suppression, necrosis induction, and inhibition of NLR-triggered responses. Further functional characterization of the effectors with their signal peptides is particularly important because many cloned resistance proteins in wheat that recognize *Z. tritici* effectors are localized in the plasma membrane. For example, Stb6 is a wall-associated kinase (WAK) with an extracellular domain that recognizes the apoplastic AvrStb6 effector (Brown et al., 2015; Saintenac et al., 2018; Zhong et al., 2017). Similarly, *Stb16q* encodes a cysteine-rich, wall-associated-like receptor kinase that likely interacts with a fungal carbohydrate to trigger resistance (Saintenac et al., 2021), and *Stb15* encodes a G-type lectin receptor-like kinase (LecRK) with extracellular domains that mediate recognition (Hafeez et al., 2025). Therefore, assessing the activity and localization of effectors with their signal peptides will provide a more comprehensive understanding of their roles in pathogen-host interactions.

In summary, we have characterized the *Z. tritici* transcriptome during susceptible, resistant, and non-host interactions and identified key temporal shifts in gene expression dynamics. A central finding of this work is that the most significant divergence in fungal transcriptional profiles between susceptible and resistant wheat cultivars occurs during the transition to necrotrophy around 10 DPI, whereas early infection stages (1 and 3 DPI) show highly similar expression patterns. In addition, comparison with the non-host interaction on barley reveals that major differences in gene expression emerge early, suggesting that non-host resistance acts during the initial stages of infection. These findings provide new insights into how *Z. tritici* tailors its transcriptional programs in response to different host environments. Pathway and gene function enrichment analyses of each interaction type identified 31 effector candidates that are upregulated early during susceptible interactions. This early induction highlights their potential roles in host colonization and immune modulation. Together, our results advance our understanding of temporal and interaction-specific transcriptional reprogramming in *Z. tritici.* Future investigations will address whether the effector candidates identified in our study have a functional role in fungal pathogenicity as well as suppress immune responses in wheat.

## Materials and Methods

### Plant material and inoculum preparation

Highly susceptible wheat cultivar Taichung 29, R-gene resistant cultivars Veranopolis (*Stb2* resistance gene) and Israel 493 (*Stb3*), and the non-host barley cultivar Kindred were grown in a greenhouse in Metro-mix 500 series potting soil (SunGro Horticulture, Bellevue, WA), with a constant 16-h light and 8-h dark cycle for 2-3.5 weeks. Wheat and barley were planted in three biological replicates consisting of 3 plants per 4-inch pot. A 1 cm^2^ piece of lyophilized filter paper stock (stored at -80°C) of isolate IPO323 of *Z. tritici* was placed into 100 ml of yeast-sucrose broth with 100 µl of kanamycin sulfate (25 mg/ml). The flask was placed on a shaker at 25°C for 3 days. The inoculum concentration was adjusted to 1.0 × 10^6^ conidia/ml, and 1 µl of Tween 20 (polysorbate 20) was added per 100 ml of inoculum. The inoculum was evenly hand sprayed over 14-days-old (2^nd^-3^rd^ leaf stage) wheat and barley plants. Inoculated plants were enclosed in a humidity chamber draped in plastic and muslin for 72 h, then removed to normal growth chamber conditions for the duration of the experiment. Emerging leaves of wheat and barley were trimmed every 2-3 days to ensure that the inoculated leaves remained “healthy” and that only inoculated tissue was harvested.

*Nicotiana benthamiana* was grown in Berger Seed Germination and Propagation Mix supplemented with Osmocote slow-release fertilizer (14-14-14). Plants were maintained in a growth chamber with a 16h:8h light:dark photoperiod at 24°C with light and 22°C in the dark and 60% humidity with average light intensities at plant height of 130 µmols/m^2^/s.

### RNA extraction and sequencing

Wheat and barley leaf tissue was harvested by clipping 2 leaves per plant per replicate and bulking 3 plants per replicate. Leaf tissue was harvested at 1, 3, 6, 10, 17 and 23 days post inoculation (DPI). All harvested leaf material was immediately frozen in liquid nitrogen. RNA was extracted using the RNeasy Plant Mini kit (Qiagen). For each replicate, 1 µg of total RNA was used to produce cDNA libraries (Purdue Genomics Core Facility, West Lafayette, IN) for RNA sequencing. The sequencing was done using Illumina HiSeq producing 100-base, paired-end (PE) reads. Read quality was checked with FastQC (Andrews, 2010) before and after trimming to remove low-quality and residual adapter sequences (Trimmomatic v.0.36). All reads were mapped to the reference genome of the *Z. tritici* IPO323 strain (*M. graminicola* v.2.0) retrieved from the Joint Genome Institute (JGI) (Goodwin et al., 2011) using BBMap (v 35.85; Bushnell B., sourceforge.net/projects/bbmap/). Reads mapped to the reference genome were counted using HTSeq (Anders et al., 2015). Fewer fungal reads were detected in all the samples of the non-host interaction, and in the samples of the resistant interactions at 17 and 23 DPI compared to the samples of the susceptible interaction, due to reduced fungal growth in the resistant wheat cultivars at the late stage and in the non-host species barley. As a result, for the comparisons of susceptible versus resistant interactions, and susceptible versus non-host interactions, we removed from the whole dataset the genes that did not have any measurable counts in the replicates of the resistant and non-host interactions at specific time points. This allowed us to improve the normalization of the counts and the consistency between replicates. This approach reduced the total number of DEGs that we were able to quantify at each time point, but we could accurately detect significant differences in the highly expressed genes in the samples and reduce the false positives. We encountered limitations in the analysis due to the low read counts obtained in the replicates of the resistant interactions and non-host interaction. These limitations include reduced sensitivity in detecting low-abundance transcripts or lower-expressed genes in the pathogen, and difficulty in assigning functional annotations or representative GO terms for some sets of DEGs.

Data were normalized and filtered using the R package DESeq2 v.4.0.3 (Love et al., 2014), which uses the median of ratios method to determine the size factors and shrinkage for variance estimation. The normalization for the comparison of susceptible versus resistant interactions was conducted on two groups of samples. One group contains the samples at 1, 3, 6 and 10 DPI, and the other group contains the samples at 17 and 23 DPI. We used this approach because very few reads were detected in the samples of the resistant interactions at 17 and 23 DPI, resulting in many genes without measurable expression. Similarly, in the comparison of susceptible and resistant interactions versus the non-host interaction, the normalization was conducted on one group that contains the samples from 1 and 3 DPI, a second group of samples from 6 and 10 DPI, and a third group that contains the samples at 17 and 23 DPI. Differential gene expression analysis for the pathogen between inoculation treatment samples within each 6-point time course was completed using DESeq2 v.4.0.3 (Love et al., 2014). Significant DEGs were determined at FDR = 5% and using a cutoff of 1-fold change in expression between comparisons.

### GO and KEGG pathway enrichment analyses

Differentially expressed genes (DEGs) were subjected to Gene Ontology (GO) enrichment analysis using gProfiler (Raudvere et al., 2019). *Mycosphaerella graminicola* v.2.0 genes with detectable counts in our analysis were used as the background reference for *Z. tritici*. The significance threshold was determined by the default gProfiler method, the g:SCS method, which corresponds to an experiment-wide threshold of α = 0.05 (Raudvere et al., 2019). KOBAS v.3.0. (Bu et al., 2021) supports *Z. tritici* and was used to calculate KEGG pathway enrichment (Kanehisa et al., 2017) of the significantly expressed genes; significance was determined by Fisher’s exact test (Fisher, 1925) using the Benjamini and Hochberg method to correct for multiple testing (Benjamini and Hochberg 1995).

### Selection of candidate effectors from *Z. tritici*

Effector predictions were performed as described previously by Helm et al. (2022) with modifications. We used SignalP (v6.0) (Teufel et al., 2022) and Phobius (v1.01) to predict signal peptide sequences and we applied a probability threshold of 0.9. The absence of transmembrane domains was predicted using Phobius (v1.01) (Käll et al., 2004) and DeepTMHMM (v1.0.24) (Hallgren et al., 2022). EffectorP (v.3.0) (Sperschneider & Dodds, 2022) was used to select *Z. tritici* proteins that contained effector-like sequence characteristics. LOCALIZER (v.1.0.4) (Sperschneider et al., 2017) and ApoplastP (v.1.0) (Sperschneider et al., 2018) were used to predict the possible localization of the candidate effectors. The list of *Z. tritici* putative effectors is shown in Table 4.

### Generation of plant expression constructs

Constructs were generated using a modified multisite Gateway cloning (Invitrogen) strategy (Helm et al., 2022; Qi et al., 2012). The 3xHA:sYFP (free super yellow fluorescent protein) construct has been described previously (Helm et al., 2022; Jaiswal et al., 2023).

The attL1 and attL4 Gateway recombination sequences were fused to the 5’ and 3’ ends of the open reading frames (ORFs) of each *Z. tritici* candidate effector, thereby generating Gate-way-compatible donor clones. A commercial gene synthesis company (Azenta Life Sciences) was used to synthesize 31 candidate effector genes without predicted signal peptides and six of the candidate effector genes with their predicted signal peptides. The sequences were inserted into the pUC57 plasmid by the service provider and designated pDONR(L1-L4):ZtEC.

To generate the ZtEC:sYFP protein fusions, the pDONR(L1-L4):ZtEC constructs were mixed with pBSDONR(L4r-L2):sYFP (Helm et al., 2022; Qi et al., 2012) and recombined into the plant expression plasmid pEG100 (Earley et al., 2006) using LR Clonase II (Invitrogen) following the manufacturer’s instructions. The resulting constructs were verified by sequencing and subsequently used for transient expression assays.

### *Agrobacterium*-mediated transient protein expression in *N. benthamiana*

*Agrobacterium*-mediated transient expression was performed as described previously (Helm et al., 2022) with modifications. The constructs described above were mobilized into *Agrobacterium tumefaciens* strain GV3101 (pMP90) and grown in Luria-Bertani (LB) broth supplemented with 25 µg of gentamicin sulfate and 50 µg of kanamycin per milliliter at 30°C for two days. Transformants were inoculated into 10 mL of liquid LB containing the appropriate antibiotics and were shaken overnight at 30°C at 250 rpm. Following overnight incubation, the cells were pelleted by centrifuging at 3,800 rpm for 3 minutes and resuspended in 10 mM MgCl_2_. The bacterial suspensions were adjusted to an optical density at 600 nm (OD_600_) of 0.5 and incubated with 100 µM acetosyringone for 3-4 hours at room temperature prior to agroinfiltration. Bacterial suspensions were infiltrated into the abaxial surface of 3- to 4-week-old *N. benthamiana* leaves with a needleless syringe.

### Laser scanning confocal microscopy

Live-cell imaging of *N. benthamiana* epidermal cells was performed forty-eight hours post agroinfiltration using a Zeiss LSM880 Axio Examiner upright confocal microscope equipped with a Plan Apochromat 20×/0.8 objective (pinhole 1.0 AU). The sYFP-tagged fusion proteins were excited using a 514-nm argon laser, and fluorescence signals were detected between 517 nm and 562 nm. All confocal micrographs were processed using the Zeiss Zen Blue Lite program. Image analysis was performed using the Fiji plugin of Image J 2.0.0 (http://fiji.sc/Fiji).

### Immunoblot analyses

Immunoblot analyses were performed as described previously with slight modifications (Helm et al., 2022; Jaiswal et al., 2023). Nine leaf discs of *N. benthamiana* (0.15 g per sample) were collected at 48 hours after agroinfiltration and flash frozen in liquid nitrogen. Frozen tissue was homogenized with 2 volumes of protein extraction buffer (150 mM NaCl, 50 mM Tris [pH 7.5], 0.1% Nonidet P-40 [Sigma-Aldrich], 1% plant protease inhibitor cocktail [Sigma-Aldrich], and 1% 2,2’-dipyridyl disulfide) using a cold ceramic mortar and pestle. The solution was centrifuged at 13,000 × g for 15 minutes at 4°C. Seventy-five microliters of the collected supernatants were combined with 25 µL of 4× Laemmli Buffer (BioRad) supplemented with beta-mercaptoethanol. Total protein samples were separated on a 4-20% Tris-glycine stain-free polyacrylamide gel (BioRad) at 180 V for 1 hour in 1× Tris/glycine/SDS running buffer. Total proteins were transferred to a nitrocellulose membrane and subsequently blocked in 5% skim milk for 1 hour at room temperature. Horseradish peroxidase-conjugated anti-GFP antibody (1:5,000) (Miltenyi Biotec) was used to detect sYFP-tagged recombinant proteins. Immunoblots were incubated for 5 minutes at room temperature with Supersignal West Femto maximum sensitivity substrates (Thermo Scientific) and developed using an ImageQuant 500 CCD imaging system (Cytiva).

## Acknowledgments

The authors thank Drs. Terri Cameron and Ian Thompson (USDA-ARS, Crop Production and Pest Control Research Unit) for technical assistance and the Purdue University Imaging Facility for access to the Zeiss LSM880 Axio Examiner upright confocal microscope. Two anonymous reviewers provided helpful comments for improving previous drafts of the manuscript. The funding body had no role in designing the experiments, collecting the data, or writing the manuscript. All opinions expressed in this paper are the authors’ and do not necessarily reflect the policies and views of USDA. USDA is an equal opportunity provider and employer.

## Funding

This research was funded by the United States Department of Agriculture, Agricultural Research Service (USDA-ARS) research project 5020-21220-014-00D.

---

**Supplementary Table 1.**
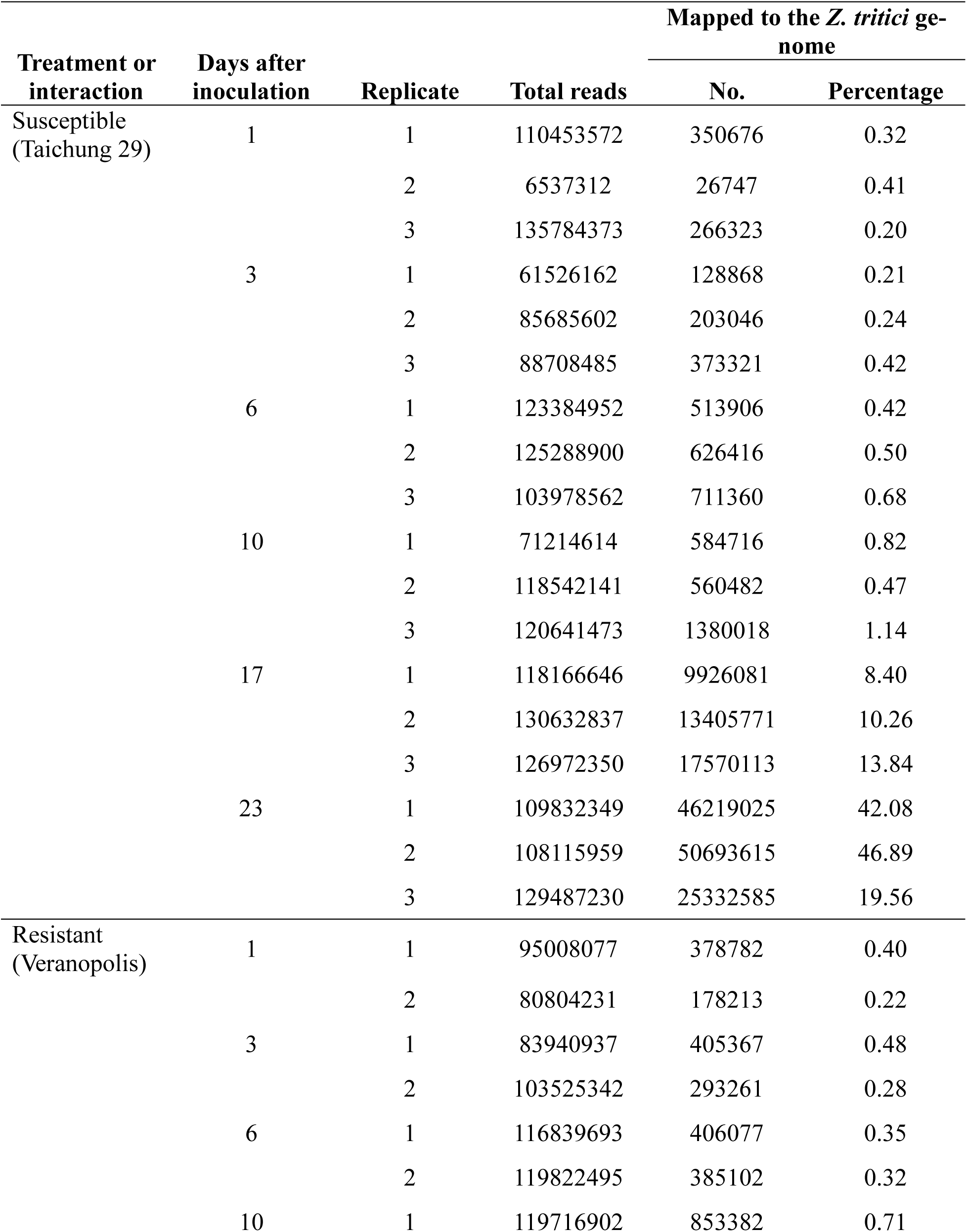
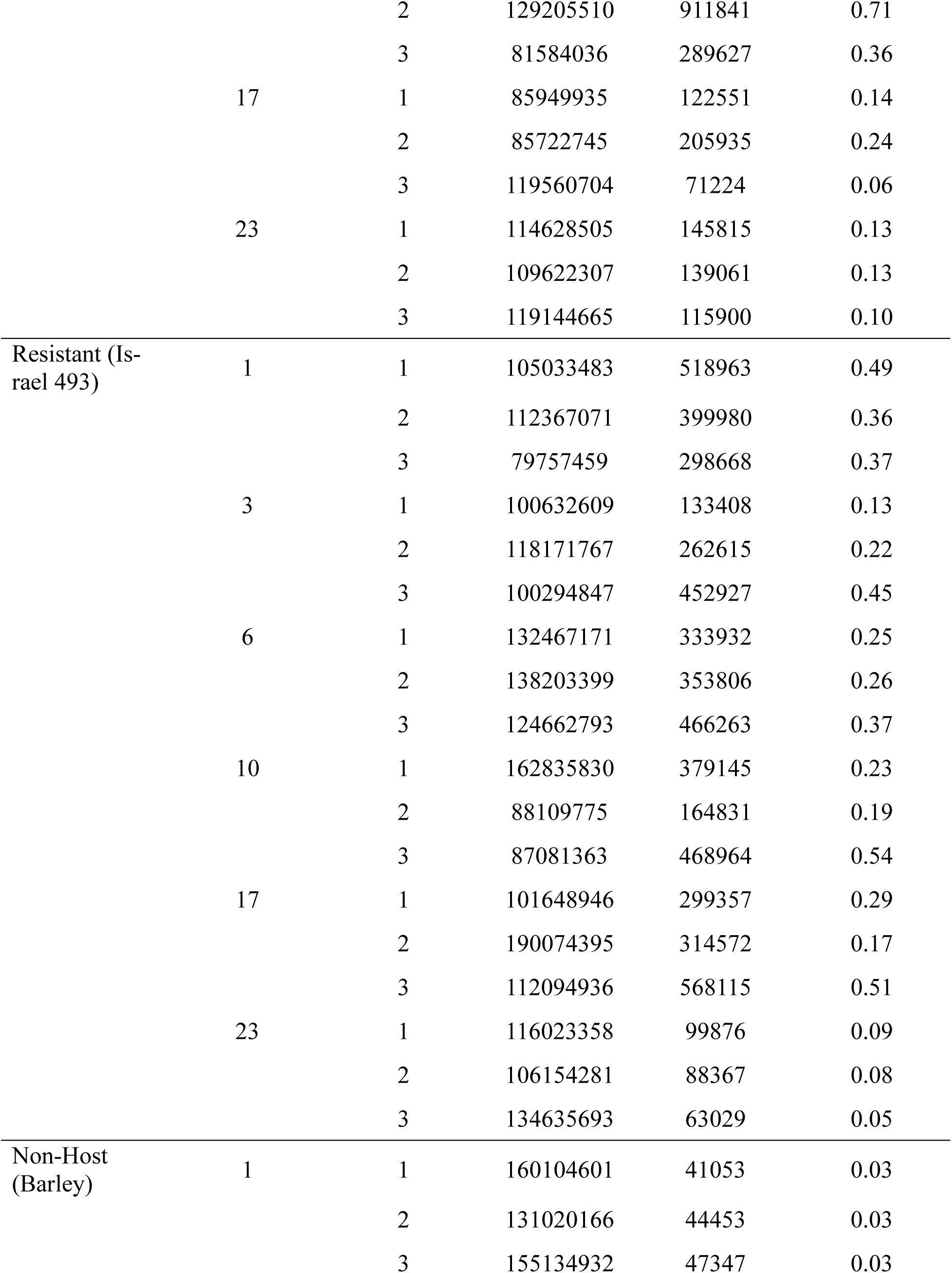
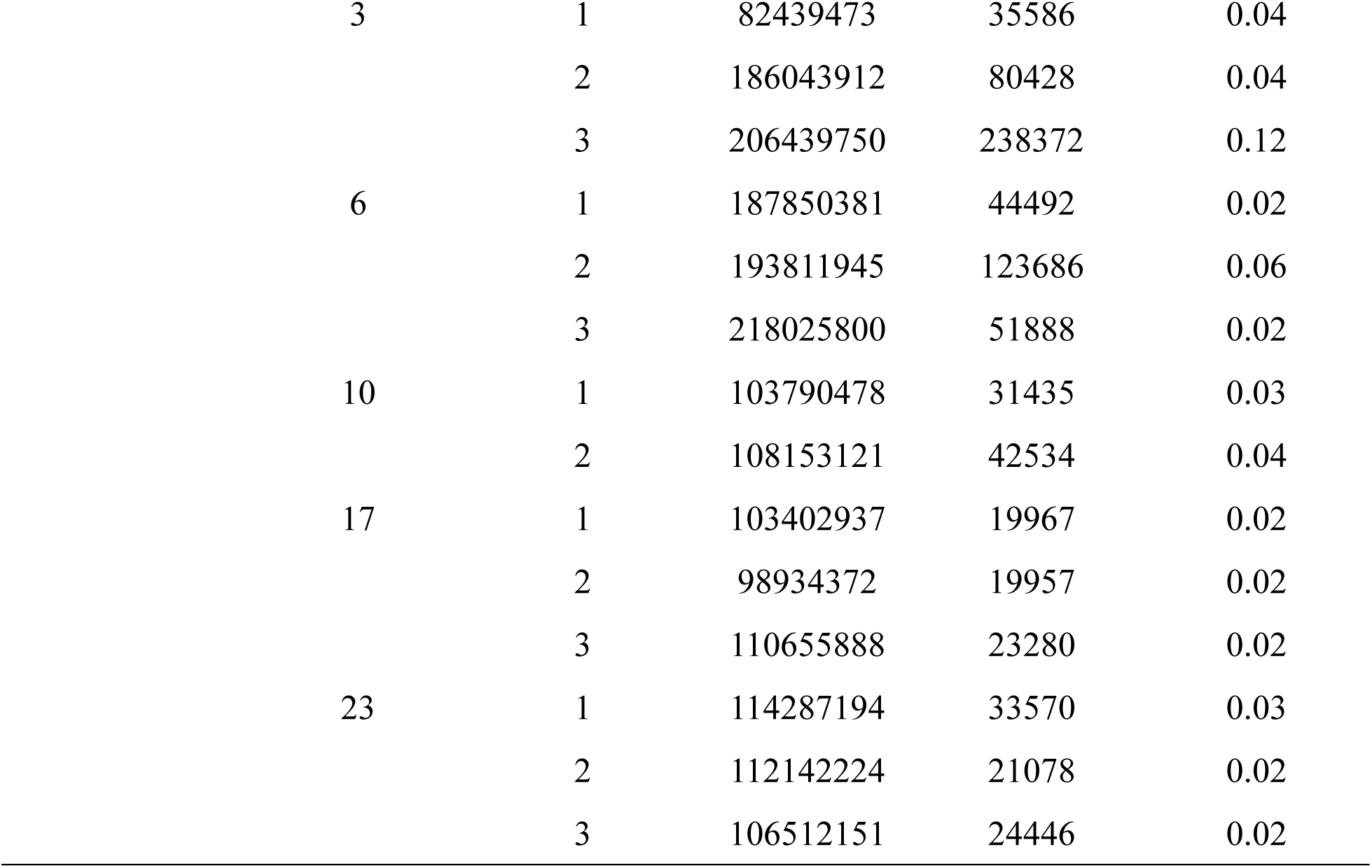
RNAseq read counts and statistics of mapped reads to the *Zymoseptoria tritici* genome.

| Treatment or interaction | Days after inoculation | Replicate | Total reads | Mapped to the *Z. tritici* genome |  |
| --- | --- | --- | --- | --- | --- |
|  |  |  |  | No. | Percentage |
| Susceptible (Taichung 29) | 1 | 1 | 110453572 | 350676 | 0.32 |
|  |  | 2 | 6537312 | 26747 | 0.41 |
|  |  | 3 | 135784373 | 266323 | 0.20 |
|  | 3 | 1 | 61526162 | 128868 | 0.21 |
|  |  | 2 | 85685602 | 203046 | 0.24 |
|  |  | 3 | 88708485 | 373321 | 0.42 |
|  | 6 | 1 | 123384952 | 513906 | 0.42 |
|  |  | 2 | 125288900 | 626416 | 0.50 |
|  |  | 3 | 103978562 | 711360 | 0.68 |
|  | 10 | 1 | 71214614 | 584716 | 0.82 |
|  |  | 2 | 118542141 | 560482 | 0.47 |
|  |  | 3 | 120641473 | 1380018 | 1.14 |
|  | 17 | 1 | 118166646 | 9926081 | 8.40 |
|  |  | 2 | 130632837 | 13405771 | 10.26 |
|  |  | 3 | 126972350 | 17570113 | 13.84 |
|  | 23 | 1 | 109832349 | 46219025 | 42.08 |
|  |  | 2 | 108115959 | 50693615 | 46.89 |
|  |  | 3 | 129487230 | 25332585 | 19.56 |
| Resistant (Veranopolis) | 1 | 1 | 95008077 | 378782 | 0.40 |
|  |  | 2 | 80804231 | 178213 | 0.22 |
|  | 3 | 1 | 83940937 | 405367 | 0.48 |
|  |  | 2 | 103525342 | 293261 | 0.28 |
|  | 6 | 1 | 116839693 | 406077 | 0.35 |
|  |  | 2 | 119822495 | 385102 | 0.32 |
|  | 10 | 1 | 119716902 | 853382 | 0.71 |
|  |  | 2 | 129205510 | 911841 | 0.71 |
|  |  | 3 | 81584036 | 289627 | 0.36 |
|  | 17 | 1 | 85949935 | 122551 | 0.14 |
|  |  | 2 | 85722745 | 205935 | 0.24 |
|  |  | 3 | 119560704 | 71224 | 0.06 |
|  | 23 | 1 | 114628505 | 145815 | 0.13 |
|  |  | 2 | 109622307 | 139061 | 0.13 |
|  |  | 3 | 119144665 | 115900 | 0.10 |
| Resistant (Is-<br>rael 493) | 1 | 1 | 105033483 | 518963 | 0.49 |
|  |  | 2 | 112367071 | 399980 | 0.36 |
|  |  | 3 | 79757459 | 298668 | 0.37 |
|  | 3 | 1 | 100632609 | 133408 | 0.13 |
|  |  | 2 | 118171767 | 262615 | 0.22 |
|  |  | 3 | 100294847 | 452927 | 0.45 |
|  | 6 | 1 | 132467171 | 333932 | 0.25 |
|  |  | 2 | 138203399 | 353806 | 0.26 |
|  |  | 3 | 124662793 | 466263 | 0.37 |
|  | 10 | 1 | 162835830 | 379145 | 0.23 |
|  |  | 2 | 88109775 | 164831 | 0.19 |
|  |  | 3 | 87081363 | 468964 | 0.54 |
|  | 17 | 1 | 101648946 | 299357 | 0.29 |
|  |  | 2 | 190074395 | 314572 | 0.17 |
|  |  | 3 | 112094936 | 568115 | 0.51 |
|  | 23 | 1 | 116023358 | 99876 | 0.09 |
|  |  | 2 | 106154281 | 88367 | 0.08 |
|  |  | 3 | 134635693 | 63029 | 0.05 |
| Non-Host<br>(Barley) | 1 | 1 | 160104601 | 41053 | 0.03 |
|  |  | 2 | 131020166 | 44453 | 0.03 |
|  |  | 3 | 155134932 | 47347 | 0.03 |

|  |  |  |  |  |
| --- | --- | --- | --- | --- |
| 3 | 1 | 82439473 | 35586 | 0.04 |
|  | 2 | 186043912 | 80428 | 0.04 |
|  | 3 | 206439750 | 238372 | 0.12 |
| 6 | 1 | 187850381 | 44492 | 0.02 |
|  | 2 | 193811945 | 123686 | 0.06 |
|  | 3 | 218025800 | 51888 | 0.02 |
| 10 | 1 | 103790478 | 31435 | 0.03 |
|  | 2 | 108153121 | 42534 | 0.04 |
| 17 | 1 | 103402937 | 19967 | 0.02 |
|  | 2 | 98934372 | 19957 | 0.02 |
|  | 3 | 110655888 | 23280 | 0.02 |
| 23 | 1 | 114287194 | 33570 | 0.03 |
|  | 2 | 112142224 | 21078 | 0.02 |
|  | 3 | 106512151 | 24446 | 0.02 |

**Supplementary Fig. S1.**
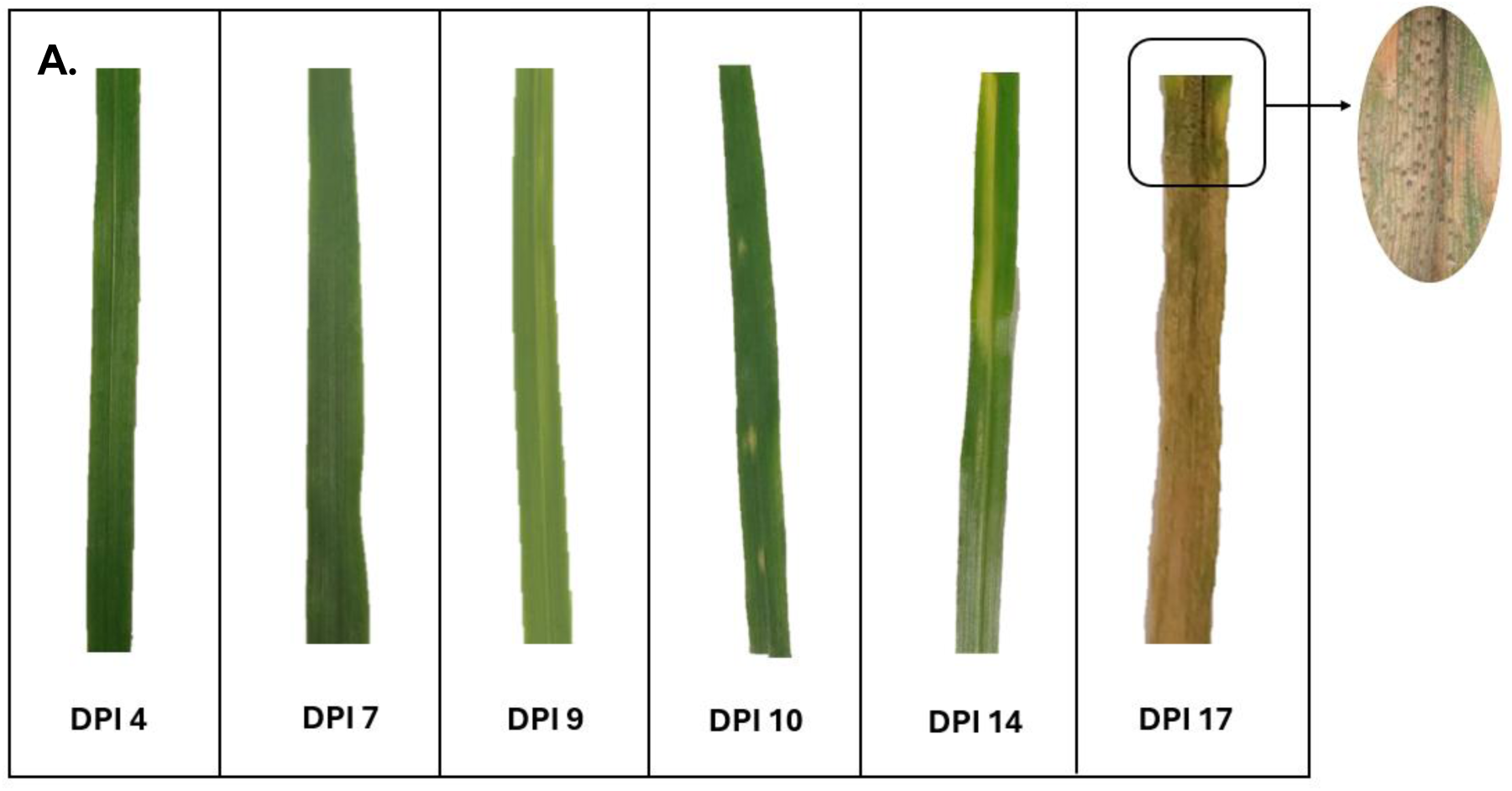
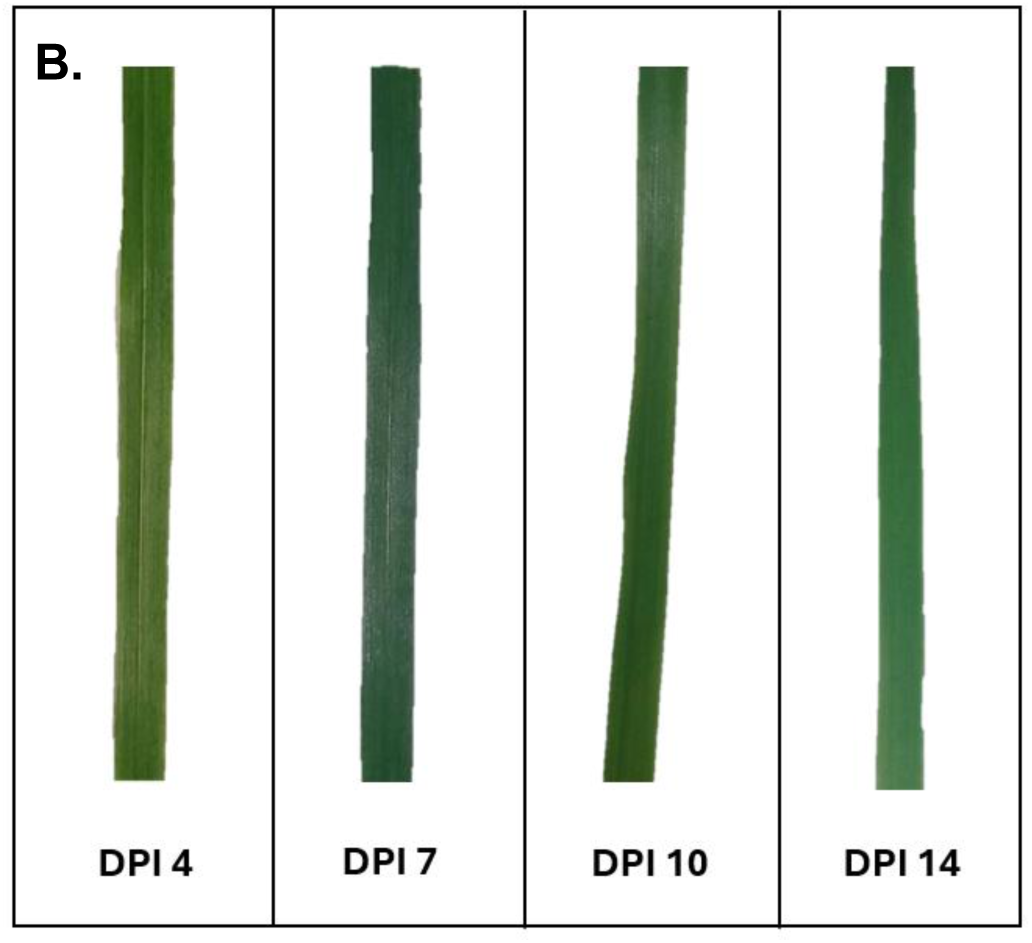
Disease establishment in the A) susceptible cultivar Taichung 29 of wheat and in the B) resistant cultivar Israel 493 at the indicated days post inoculation (DPI) with the wheat pathogen *Zymoseptoria tritici*. A closeup of the susceptible response at 17 DPI is shown in the expanded view at the upper right.

**Supplementary Fig. S2.**
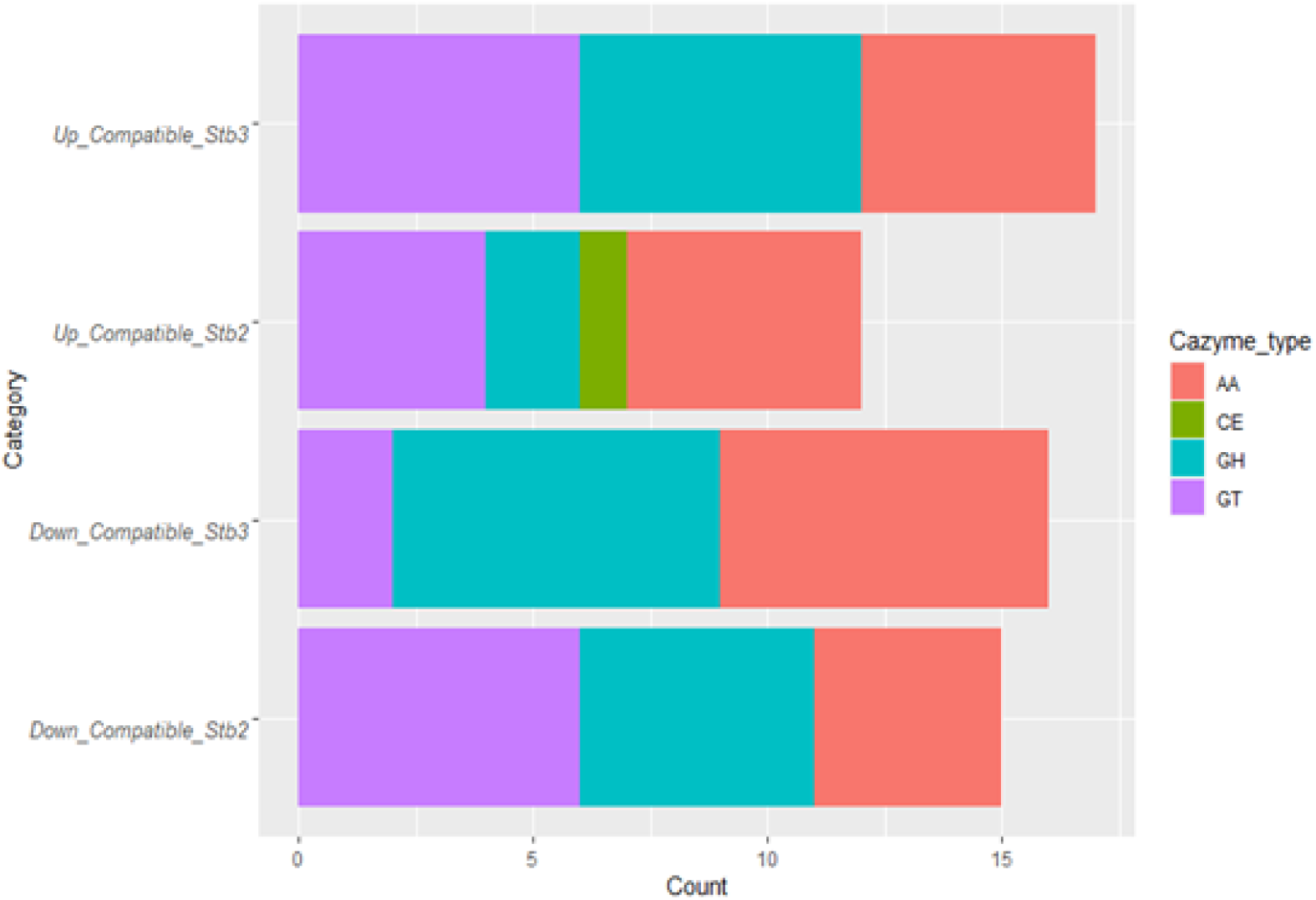
Carbohydrate-active enzymes of *Zymoseptoria tritici* identified at 10 days post inoculation in the comparison between susceptible and resistant interactions. Compatible = Susceptible.

**Supplementary Fig. S3.**
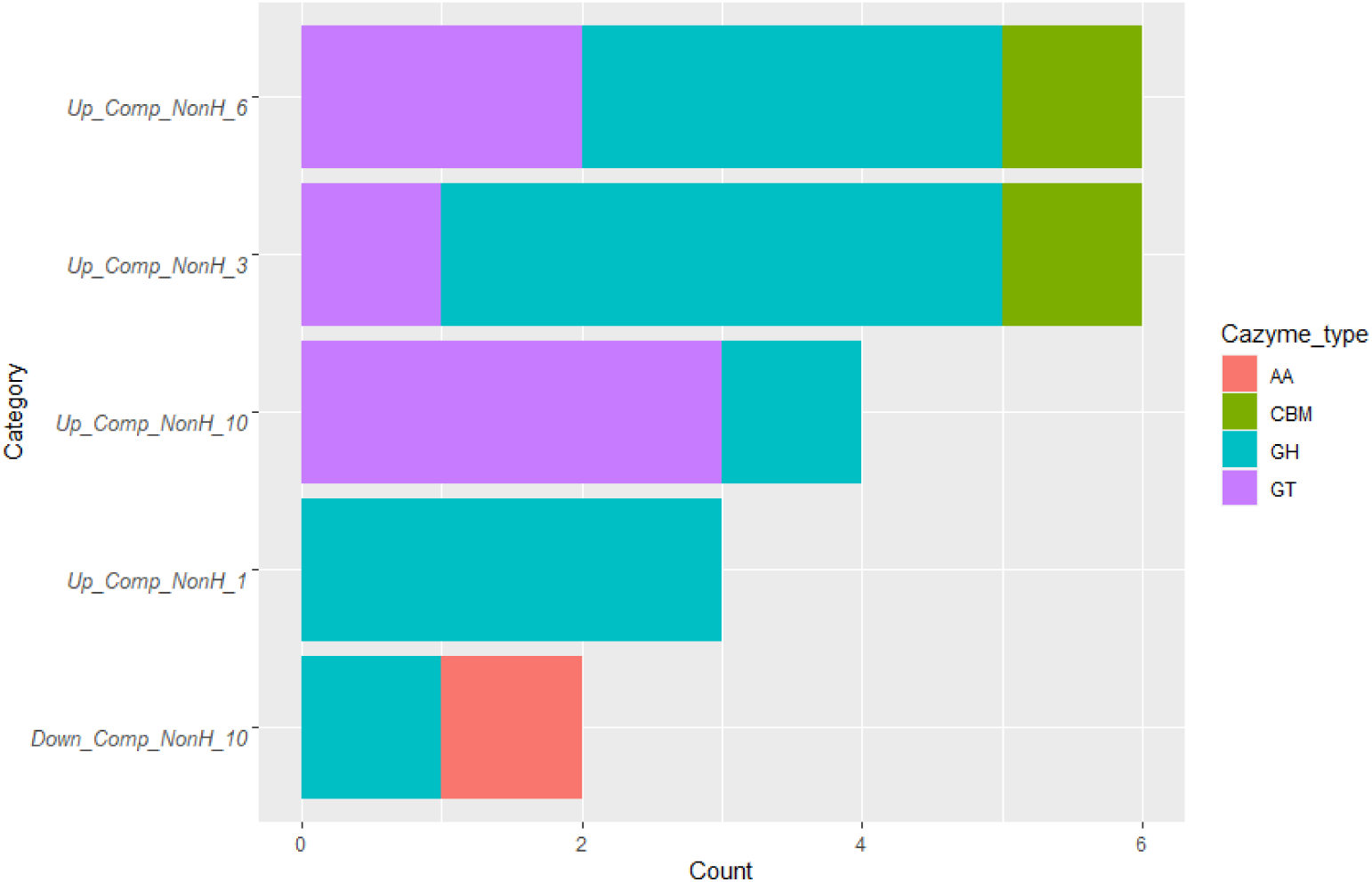
Carbohydrate-active enzymes of *Zymoseptoria tritici* identified at 6, 3, 10, 1 and 10 days post inoculation (from top to bottom) in the comparison between susceptible and non-host interactions. Comp = Susceptible, NonH= Non-Host.

**Supplementary Fig. S4.**
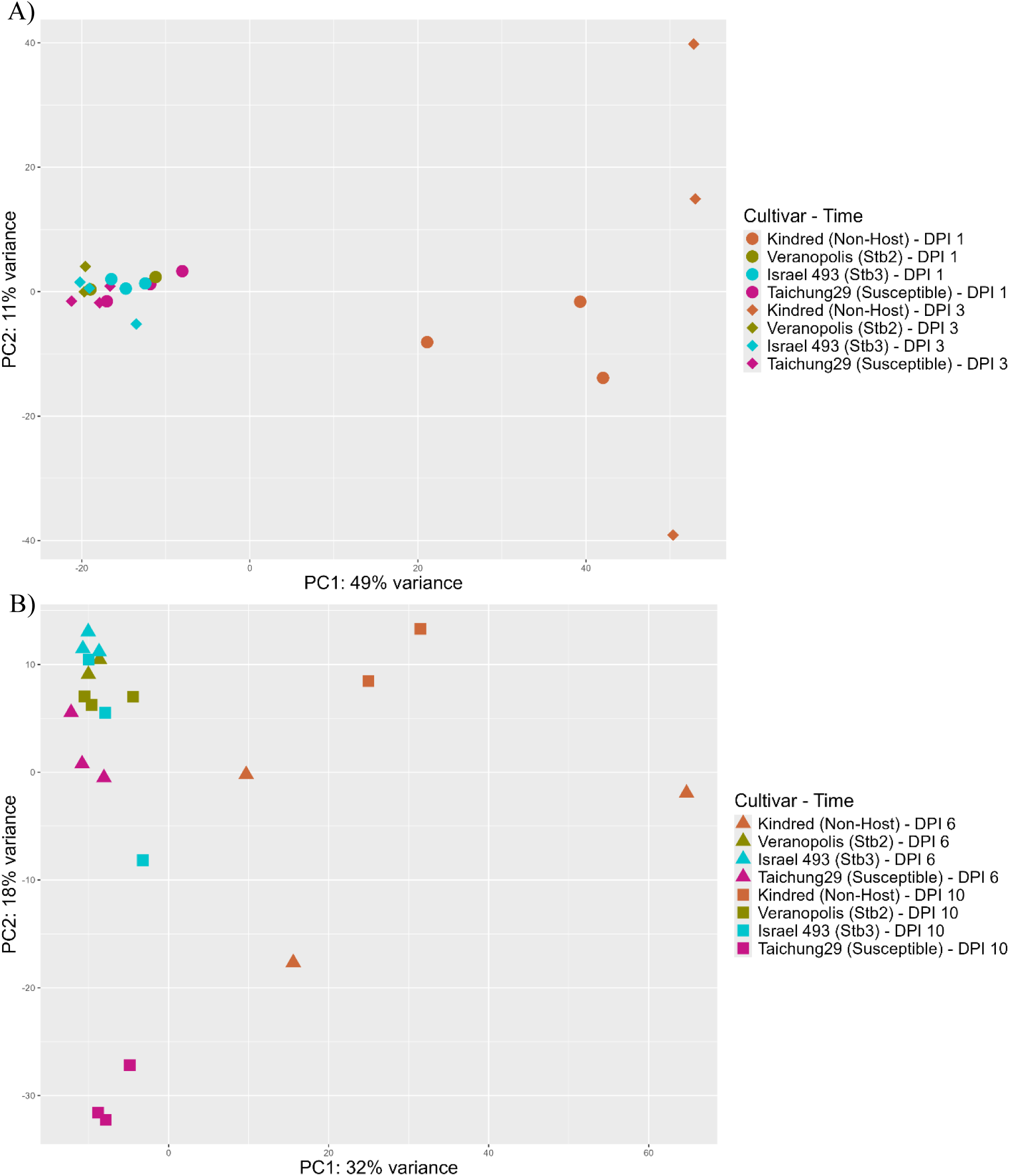
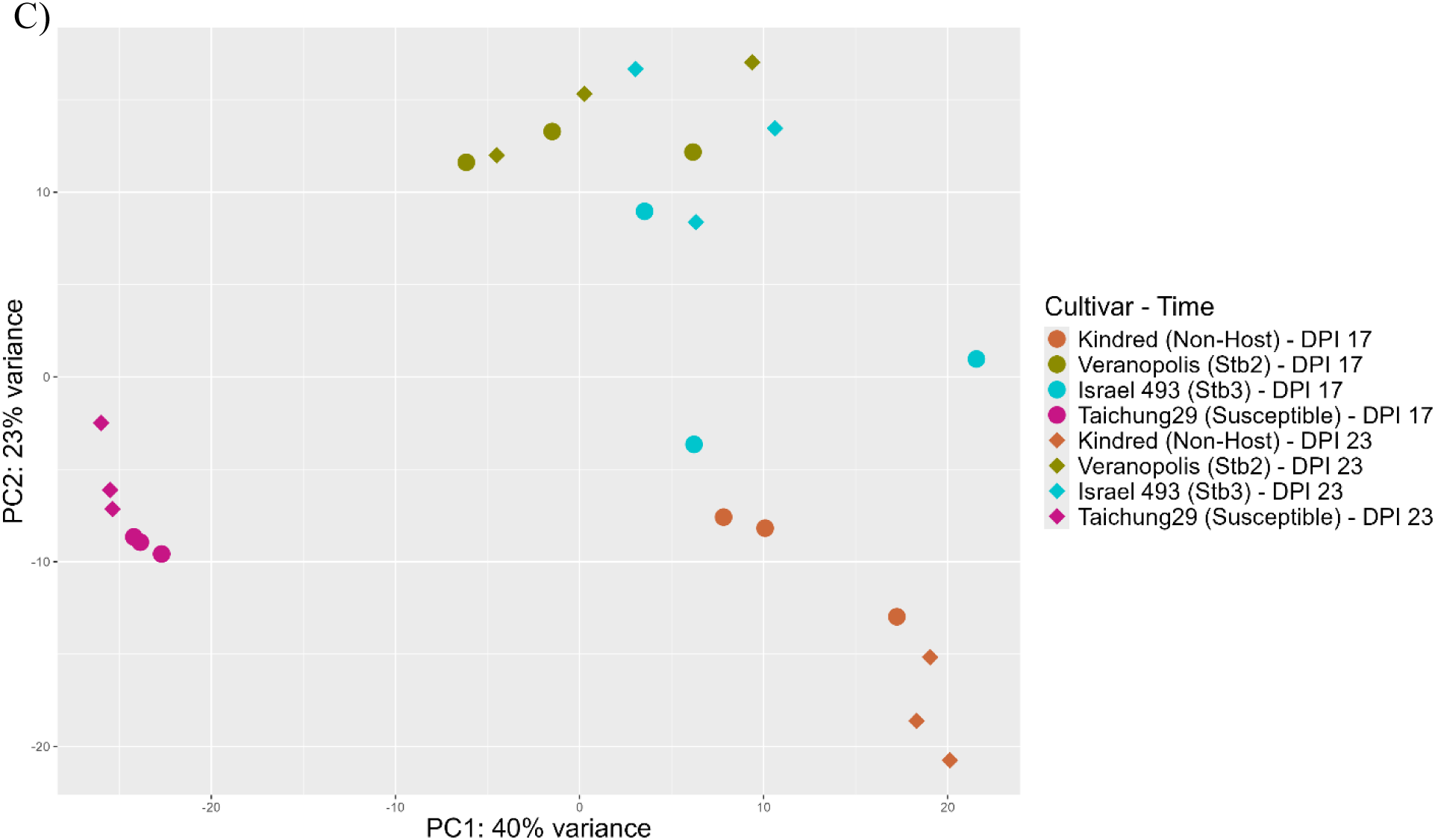
Principal component analysis (PCA) of the normalized filtered read counts for RNA of *Zymoseptoria tritici* inoculated onto the highly susceptible wheat cultivar Taichung 29 (Susceptible) or the R-gene-containing resistant wheat cultivars Veranopolis (*Stb2*) or Israel 493 (*Stb3*) or the non-host barley at: A) 1 and 3; B) 6 and 10; and C) 17 and 23 days post inoculation (DPI).

**Supplementary Fig. S5.**
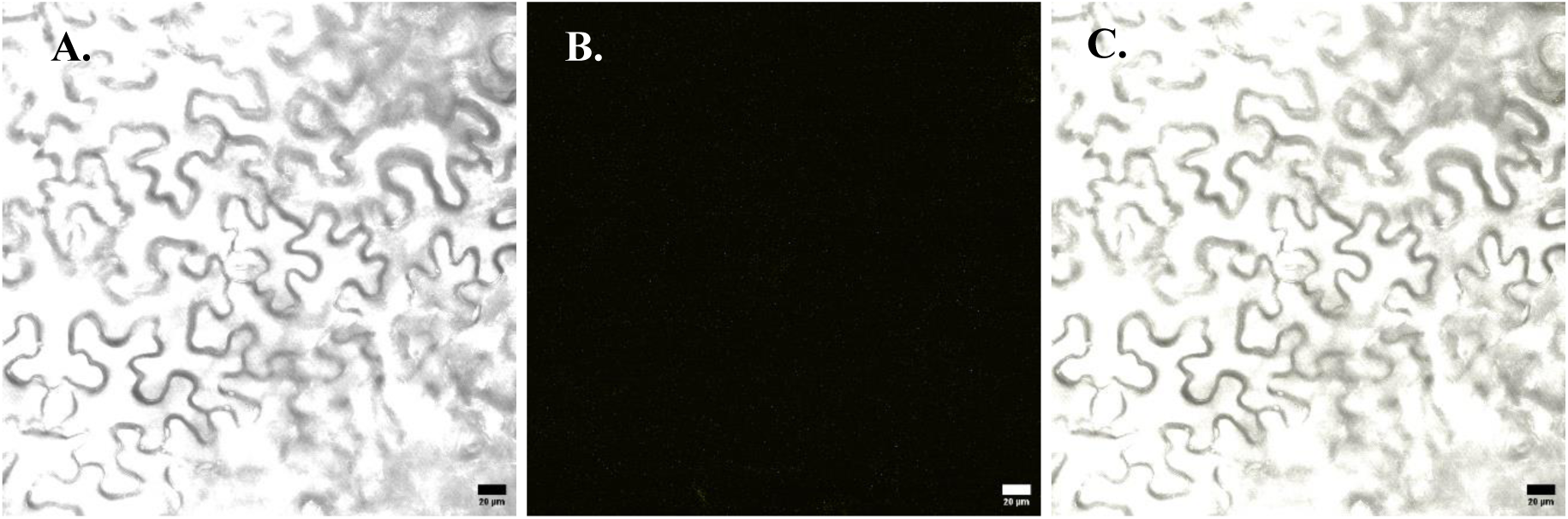
Image of a *Nicotiana benthamiana* leaf infiltrated with *Agrobacterium* expressing a protein with an -HA tag showing: A) brightfield; B) wavelength used for sYFP (514 nm) excitation; and C) the composite image. The gain was increased on the confocal microscope to enhance the detection of any potential low-level fluorescence observed in the leaf tissue.

**Supplementary Fig. S6.**
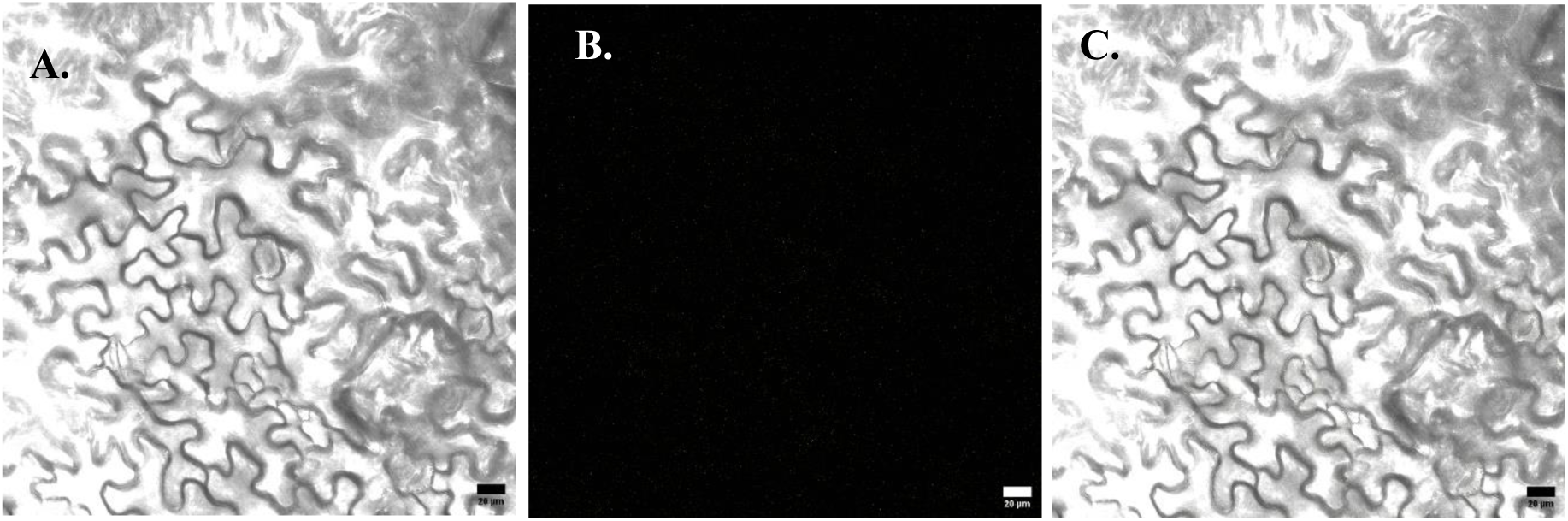
Image of a non-infiltrated *Nicotiana benthamiana* leaf showing: A) brightfield; B) wavelength used for sYFP (514 nm) excitation; and C) the composite image. The gain was increased on the confocal microscope to enhance the detection of any potential low-level autofluorescence in the leaf tissue.

**Supplementary Fig. S7.**
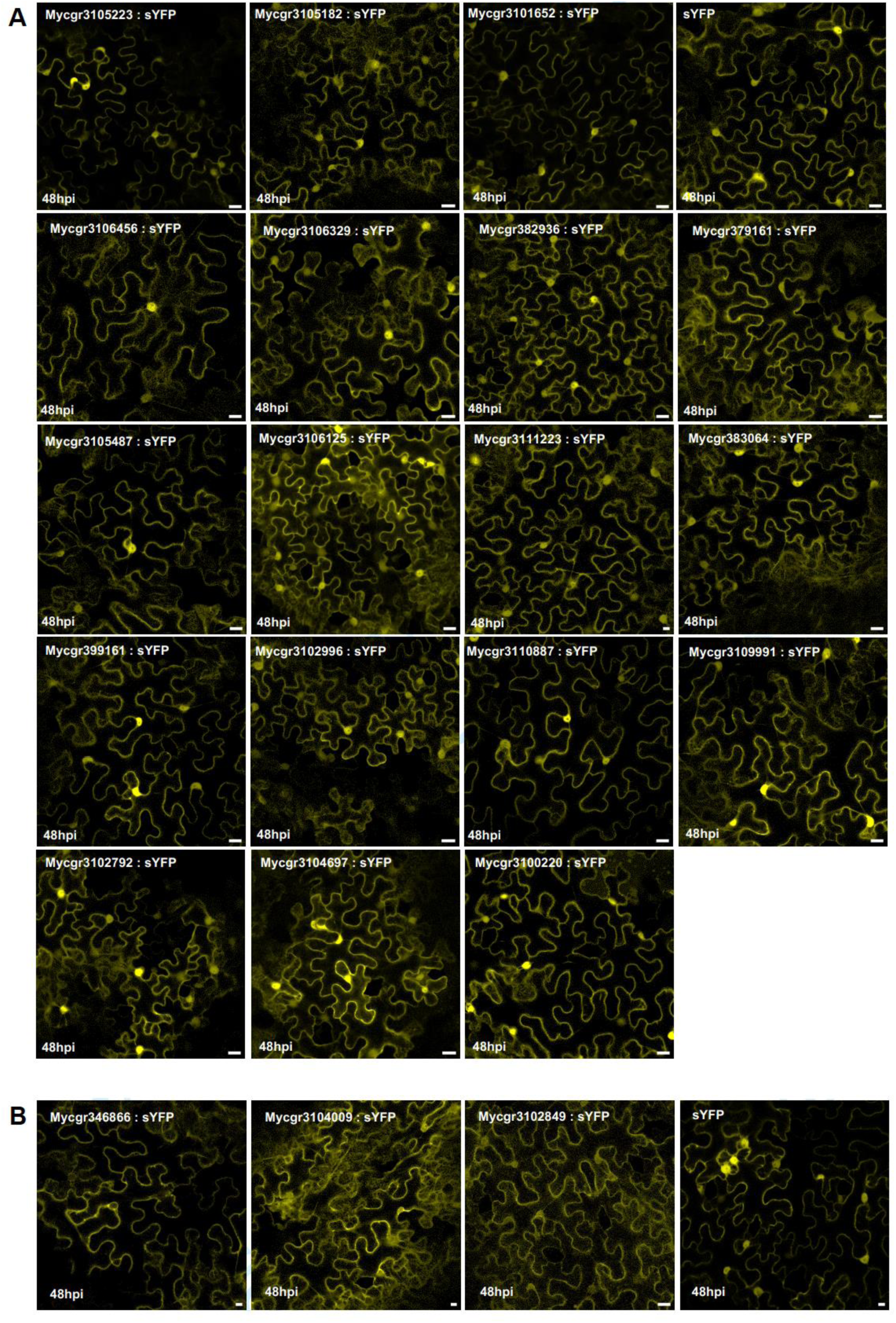
Subcellular localization of *Zymoseptoria tritici* effectors in *Nicotiana benthamiana* leaf cells expressed without their putative signal peptides. **A)** Z*ymoseptoria tritici* candidate effectors display a non-informative nucleus-cytosol localization in *N. benthamiana* that is indistinguishable from the localization of the free sYFP control. **B)** Four *Z. tritici* candidate effectors localize predominantly to the cytosol. sYFP-tagged *Z. tritici* candidate effectors were transiently expressed without their putative signal peptides in *N. benthamiana* leaves using agroinfiltration. Confocal micrographs of leaf epidermal cells were captured 48 hours following agroinfiltration. The scale bars shown represent 20 µm. All confocal micrographs are of single optical sections.

**Supplementary Fig. S8.**
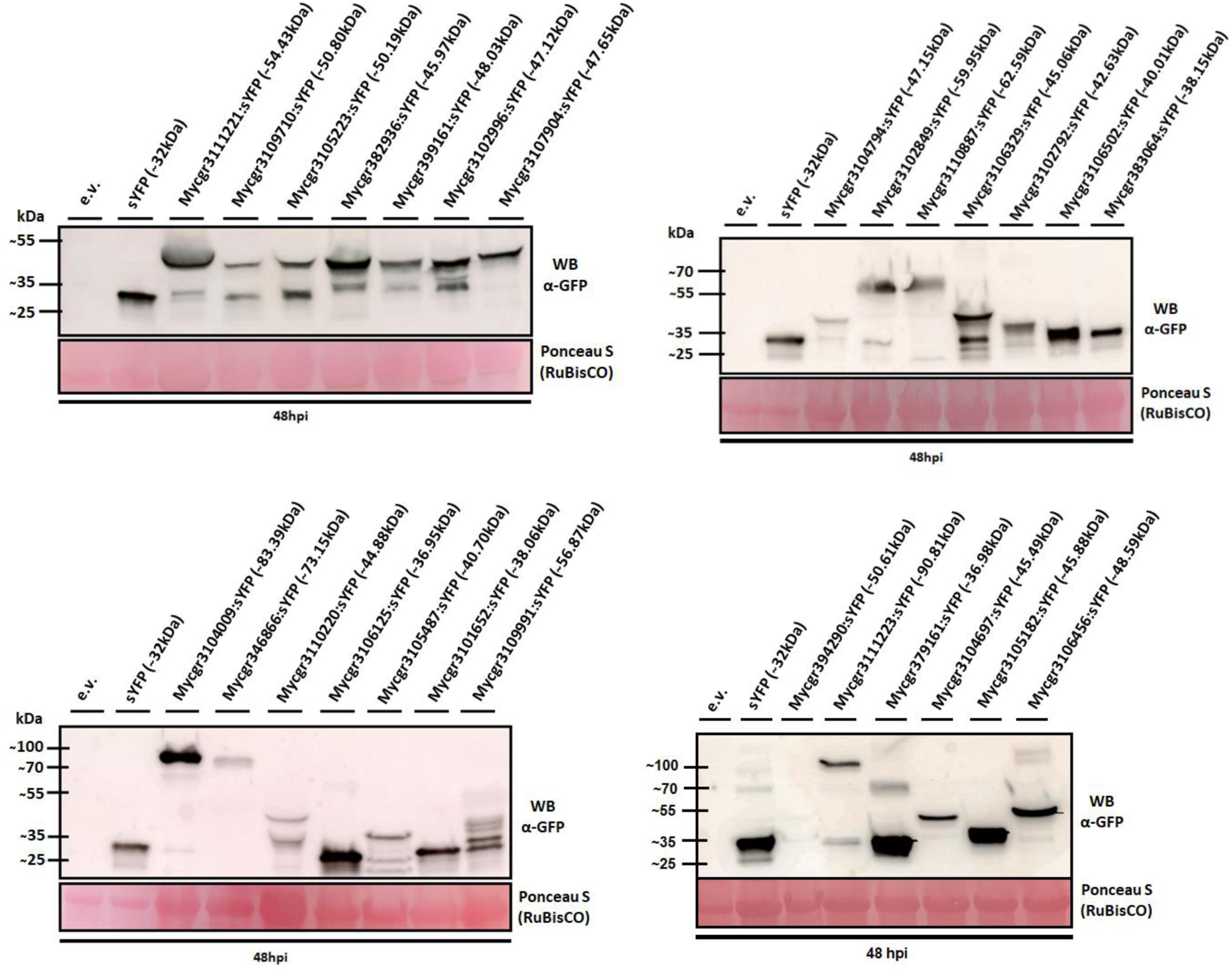
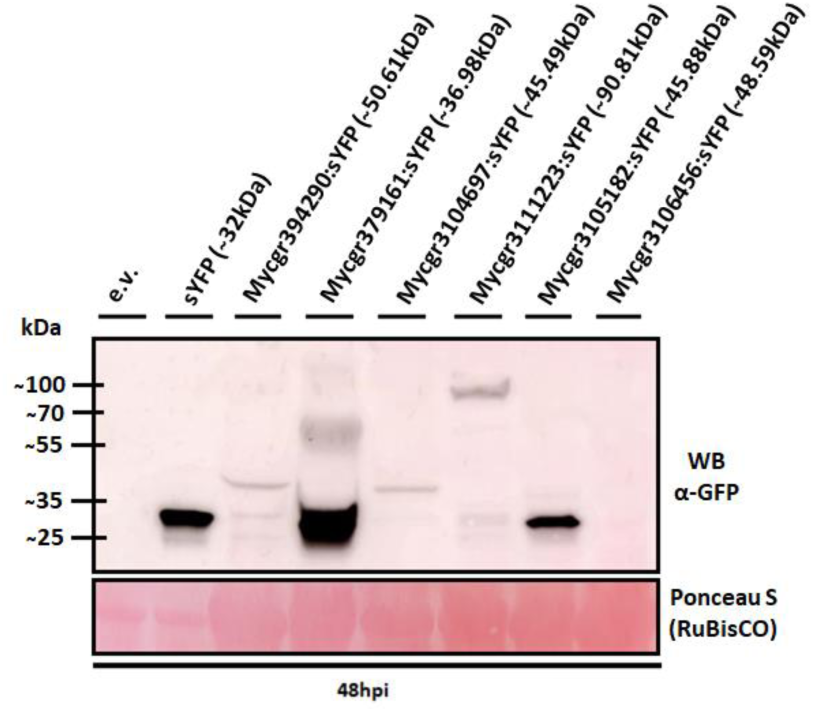
Immunoblot analyses of the *Zymoseptoria tritici* effector-fluorescent protein fusions expressed without their signal peptides. Ponceau staining of the large subunit of RuBisCO served as a loading control.

**Supplementary Fig. S9.**
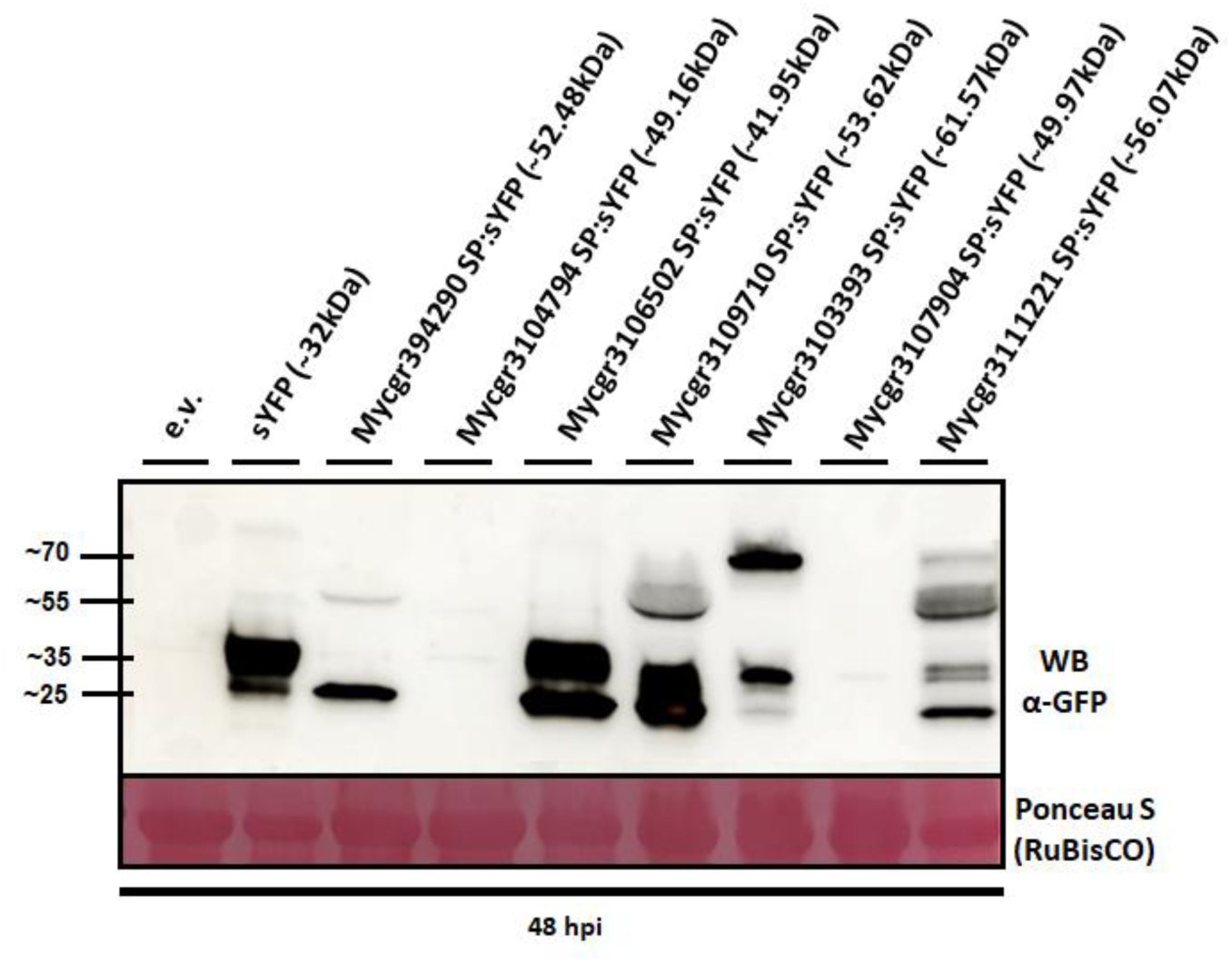
Immunoblot of the *Zymoseptoria tritici* effector-fluorescent protein fusions expressed with their signal peptides. Ponceau staining of the large subunit of RuBisCO served as a loading control. Candidate effector Mycgr3103393 was not included in the selected set of thirty-one candidate effectors, but it is shown in this immunoblot. This figure was previously published in “Functional characterization of *Zymoseptoria tritici* candidate effectors reveals their role in modulating immunity in *Nicotiana benthamiana*” by S. V. Gomez-Gutierrez et al., 2025 (bioRxiv preprint: https://doi.org/10.1101/2025.05.20.655180). Reprinted with permission.

## Notes

### Competing Interest Statement

The authors have declared no competing interest.

### Summary of Updates

Title and other changes were made in response to anonymous reviewers.

